# Single-crypt evolutionary trajectories of human sporadic and hereditary colorectal precancers

**DOI:** 10.64898/2026.04.13.718204

**Authors:** Shanlan Mo, Duo Xie, Hao Zhang, Kun Wang, Juanzhen Tong, Li Fan, Daqi Deng, Jiancong Hu, Jing Yu, Zan Li, Leo T.O. Lee, Weiwei Zhai, Zhen He, Zheng Hu

## Abstract

Our understanding of the earliest genomic processes and evolutionary trajectories that distinguish sporadic from hereditary human cancers remains limited, impeding early diagnosis and risk stratification. Here, we performed whole-genome sequencing of 319 single crypts dissected from normal colorectal mucosa, premalignant polyps and malignant colorectal cancers (CRCs) from five sporadic patients and three familial adenomatous polyposis (FAP) patients. We found 66% of premalignant crypts from sporadic polyps did not harbor *APC* mutations, but frequently accumulated non-canonical but positively selected driver mutations including *KMT2C, CACNA1A and FBLN2*. In contrast, most premalignant crypts (71%) in FAP indeed acquired one or more somatic *APC* mutations or 5q loss of heterogeneity (LOH). Sporadic polyps were dominated by clock-like mutational processes at single-crypt level. In contrast, FAP crypts evolved more diverse genomics processes, including reactive oxygen species-associated damages, APOBEC-associated mutagenesis and *pks*^*+*^ *Escherichia coli-*associated mutagenesis. Premalignant crypts in FAP also exhibited a 2-fold higher burden of LINE-1 retrotranspositions than sporadic ones, despite arising at a substantially younger age. Phylogenetic timing estimations showed that the first somatic *APC* mutation in FAP often occurs prenatally. Phylogeographic analysis further revealed spatial lineage segregation in early polyclonal polyps, followed by Big-Bang lineage intermixing in later monoclonal ones. Together, these data demonstrate that early colorectal tumorigenesis does not obey a universal *APC*-first model, where sporadic and hereditary precancers follow distinct evolutionary trajectories. Our study also highlights that polyclonal-to-monoclonal transition in premalignant stages represents the Big-Bang growth of colorectal tumors, offering crucial insight into early diagnosis and risk stratification.

## Introduction

Understanding the earliest evolutionary steps of tumorigenesis is crucial for cancer risk stratification and prevention^1-6^. Unlike malignant cancers, which are evolutionarily late in tumorigenesis^7,8^, premalignant lesions provide natural “fossils” recording early evolutionary processes^5,9,10^. Colorectal cancers (CRCs) develop through the canonical adenoma-carcinoma sequence, occurring in both sporadic or hereditary forms^11^. Sporadic CRCs accounts for more than 90% of cases and is generally thought to emerge through the gradual accumulation of somatic alterations in canonical CRC driver genes (such as *APC, KRAS, TP53*), leading to the malignant transformation from premalignant adenoma or polyps^12,13^. Hereditary CRC develops in a background of germline predisposition. Familial adenomatous polyposis (FAP), usually caused by a germline *APC* mutation, accounts for ∼1% of all CRCs characterized by the early appearance of hundreds to thousands of adenomatous polyps throughout the colorectum^14,15^. Without prophylactic intervention, FAP patients are at an extremely high-risk of developing CRC. Therefore, examining sporadic and FAP premalignant polyps provide critical insights into the tempo and mode of tumor initiation and evolution^16-19^.

Through lineage tracing combined with genomic analysis, we and others have demonstrated that early intestinal lesions are commonly of polyclonal origins in both mouse models^19-22^ and human premalignant polyps^18,19^. These findings challenge the classical view that a colorectal neoplasia is initiated by a single progenitor cell^23^. Instead, early lesions may arise through cooperation among multiple progenitor clones before later monoclonal progression^18-20,24^. However, a systematic characterization of genomic landscape and evolutionary dynamics that distinguish sporadic from hereditary precancers are still unclear. This is crucial for refining risk stratification and developing interception strategies tailored to distinct evolutionary trajectories.

The normal intestinal epithelium and neoplasia are both organized into glandular units known as crypts, or glands. Each human crypt contains several thousands of cells^25^ and is maintained by a small number of stem cells in normal conditions^26^. This glandular spatial architecture in the intestine enables the sequencing of individual crypts as naturally occurring clones, resolving somatic clonal relationships at near single-clone resolution. Although single-crypt sequencing has been widely used to study the normal colorectal epithelium^27,28^, inflammatory bowel disease (IBD)^29^ and CRC^30-32^, its use in premalignant polyps remains limited. Consequently, the genomic processes and evolutionary dynamics operating in premalignant polyps at single-clone resolution are still poorly understood.

To fill this gap, we generated a large-scale, multi-regional single-crypt whole-genome sequencing (WGS) datasets to define the genomic trajectories of human sporadic and hereditary colorectal polyps. We profiled a total of 46 tissues (or lesions) spanning normal mucosa, premalignant polyps and synchronous tumors from five sporadic patients and three FAP patients. A validation cohort of 107 colorectal polyps with tissue-level bulk whole-exome sequencing (WES) was also included^19^. Through a comprehensive analysis of somatic mutations, copy-number alterations, LINE-1 retrotransposition and phylogenetic trees, we identified distinct genomic and evolutionary processes operating in sporadic and FAP tissues. We further show that the polyclonal-to-monoclonal transition is accompanied by a spatial shift from lineage segregation to lineage intermixing, which is consistent with the onset of Big-Bang tumor growth.

## Results

### Single-crypt WGS of synchronous colorectal polyps and cancers

We collected freshly resected specimens from five sporadic patients with synchronous CRC and polyps and three FAP patients. Two of the three FAP patients had developed CRCs by the time of surgery, at ages 35 (FAP01) and 24 (FAP04) (**Supplementary Table 1**). None of the sporadic patients had a history of inflammatory bowel disease (IBD). In total, 46 tissue samples were collected, spanning from normal mucosa (n = 16) and premalignant polyps (n = 26) to synchronous malignant adenocarcinomas (n = 4) (**Fig. 1a, Supplementary Fig. 1, Supplementary Tables 1-2**).

**Figure 1.**
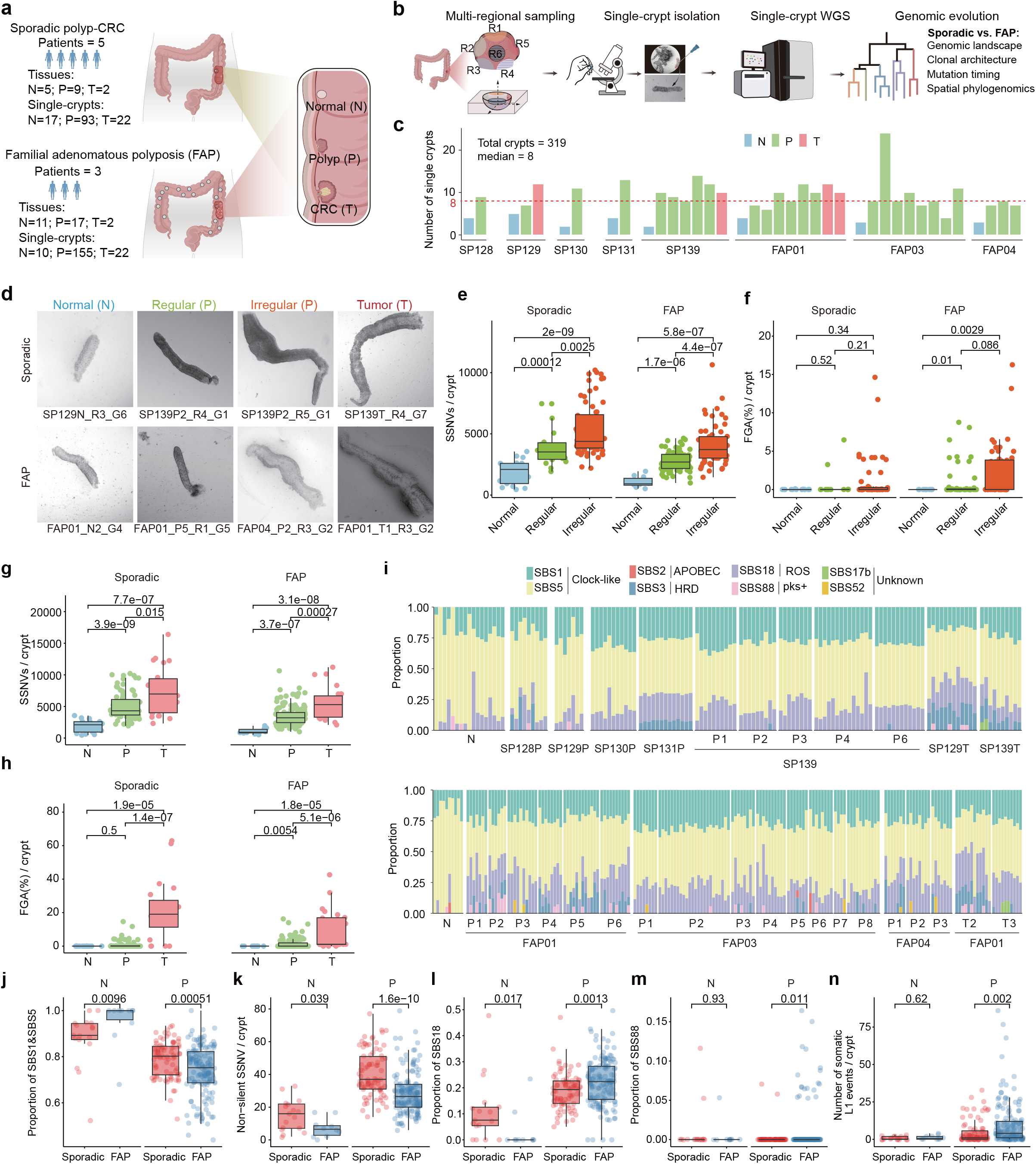
Single-crypt WGS reveals differential genomic processes in sporadic and hereditary colorectal precancers. **a**, Overview of the human single-crypt WGS including five sporadic patients and three FAPs. **b**, Schematic of single-crypt WGS procedure. The single crypts from multiple regions of each tissue were isolated under microscope and were subjected to WGS. **c**, Number of sequenced crypts from each tissue per patient. **d**, Morphological characterization of individual crypts spanning from normal mucosae, premalignant polyps and synchronous tumors. **e**, Comparison of the somatic mutation burden among normal crypts, regular and irregular polypous crypts from sporadic (left) and FAP (right) patients. **f**, Comparison of FGA across normal crypts, regular and irregular polypous crypts from sporadic (left) and FAP (right) patients. **g**, Somatic mutation burden per crypt across normal (N), polyp (P) and tumor (T) samples from sporadic (left) and FAP (right) patients. **h**, The fraction of genome altered (FGA) per crypt across N, P and T samples from sporadic (left) and FAP (right) patients. **i**, The proportional contribution of single-base-substitution (SBS) signatures in each crypt from sporadic (top) and FAP (bottom) patients. Single crypts are grouped by corresponding patients, with crypts from normal tissue, polyps and tumors shown separately. Signatures matched with potential molecular processes are color-coded as indicated at the top. **j**, The proportion of SBS1 and SBS5 mutations in normal and polypous crypts from sporadic and FAP patients. **k**, The number of non-silent somatic mutations in normal and polypous crypts from sporadic and FAP. **l**. The proportion of SBS18 mutations in normal and polypous crypts from sporadic and FAP. **m**. The proportion of SBS88 mutations in normal and polypous crypts from sporadic and FAP. **n**. The number of LINE-1 events in normal and polypous crypts from sporadic and FAP. Box plot horizontal line denotes the median and the box denotes the 25th to 75th percentiles, and the whiskers extend to 1.5 times the interquartile range. *P*-value, two-sided Wilcoxon rank-sum test. The colon in **a** and **b** was adapted from the licensed images (License ID:OAY-YP4f3bb) provided by Figdraw (www.figdraw.com).

Single crypts were isolated following established protocols^19,30^, with microscopic inspection used to verify glandular morphology (**Fig. 1a-c**). Multiple spatial regions (∼5 per tissue) were dissected from each tissue. Individual crypts were then isolated (∼8 crypts per tissue**; Fig. 1c**) and whole-genome sequenced at a mean depth of ∼24X using a low-input DNA library preparation protocol^33^ (**Supplementary Table 3**). Somatic single nucleotide variants (SSNVs), short insertions/deletions (indels), and somatic copy number alterations (SCNAs) were identified using in-house bioinformatic pipelines as our published studies^19,34,35^ (**Methods**).

Among the 248 polypous crypts examined, 138 (55.6%) showed an irregular phenotype characterized by abnormal morphology and increased length (**Fig. 1d, Supplementary Fig. 2, Methods**). The remaining 38.7% retained a relatively regular morphology, despite harboring significantly more SSNVs than normal mucosal crypts (**Fig. 1e**). Notably, irregular crypts generally harbored significantly more SSNVs than morphologically regular ones (**Fig. 1e**), indicating that genetic and morphological changes evolve progressively during precancerous growth (**Supplementary Fig. 2g**). Polypous crypts from both sporadic and FAP patients showed low levels of SCNAs (mean fraction of genome altered or FGA: 1.32%), though irregular crypts in FAP showed relatively higher SCNAs burdens (**Fig. 1f**). Consistent with this, SSNV burden in single crypts increased from normal tissues to polyp and then to tumors, whereas FGA remained low in most premalignant crypts (**Fig. 1g-h, Supplementary Fig. 3**).

These findings indicate that morphologically regular crypts in premalignant polyps have already accumulated substantial somatic mutations, as 2-3 folds of adjacent normal crypts (∼2.1-fold in sporadic and ∼2.6-fold in FAP). Therefore, continuous mutation accumulation precedes morphological changes characterizes the earliest stage of CRC initiation.

### Distinct genomic signatures in sporadic and FAP polyps

Somatic mutational signatures reflect the activity of endogenous cellular processes, exposure to exogenous environmental or lifestyle mutagens^36^. To evaluate the mutational activity at the single-crypt level, we extracted the single base substitutions (SBS) signatures using a hierarchical Dirichlet process (HDP)^37^, and estimated the mutational processes of each crypt using sigfit^38^ (**Fig. 1i, Supplementary Table 3**). We identified eight signatures showing strong similarity to reference signatures in the Catalogue of Somatic Mutations in Cancer (COSMIC) database, SBS1 and SBS5 as clock-like signatures, SBS2 caused by catalytic polypeptide-like (APOBEC) DNA-editing enzymes^39^, SBS3 caused by homologous repair deficiency, SBS18 caused by DNA damage by reactive oxygen species (ROS)^40^, SBS88 caused by *pks*^*+*^ *Escherichia coli (E*.*coli)*^41^, SBS17b and SBS52 with unknown cause (**Fig. 1i**).

Somatic mutations in both normal and polypous crypts were dominated by clock-like signatures (SBS1 and SBS5) (**Fig. 1j**). This pattern was more pronounced in sporadic polyps than in FAP polyps, consistent with the older ages (mean age 64 of sporadic vs. 32 years old of FAP) and more non-silent mutations in sporadic patients than in FAP patients (**Fig. 1k**). By contrast, FAP polyps showed significantly higher ROS-associated SBS18 mutations (**Fig. 1l**) and *pks*^*+*^ *E*.*coli*-associated SBS88 mutations (**Fig. 1m**). In addition, SBS2, probably caused by APOBEC family cytidine deaminases, was only detected in FAP crypts (**Fig. 1i**). These data highlight that sporadic colorectal tissues are mainly driven by clock-like mutational processes, whereas FAP tissues evolve through a more diverse mutational process.

LINE-1 (L1) retrotransposition is increasingly recognized as a source of genomic instability in cancer^42,43^, as exemplified by CRC^43^ and normal colorectal epithelium^44^. Using two complementary bioinformatic approaches, xTea^45^ and DELLY^46^, we identified 2791 somatic L1 insertions in total in our single-crypt WGS data (**Methods**). Interestingly, FAP polyps exhibited significantly higher L1 retrotransposition activity than sporadic polyps (**Fig. 1n**), suggesting that L1 mobilization accompanies to the early clonal expansion of FAP tissues. Together, these results further support that hereditary and sporadic colorectal tissues evolve through distinct genomic processes.

### Distinct CRC drivers underlie sporadic and hereditary polyp initiation

Having observed marked differences in genomic processes, we next asked whether sporadic and hereditary polyps also differ in their driver mutation landscapes at the single-crypt level (**Fig. 2, Supplementary Fig. 4**). Somatic *APC* mutations were rare in normal mucosal crypts (10% or 1 out of 10 in FAP, 6% or 1 out of 17 in sporadic patients), consistent with previous study^18,27,29,47^. Unexpectedly, only 34% of premalignant crypts from sporadic polyps harbored somatic *APC* mutations (**Fig. 2a-b, Supplementary Fig. 4a**). In contrast, 71% of premalignant crypts from FAP indeed carried one or more somatic *APC* mutations or 5q LOH, in addition to the germline *APC* mutation (**Fig. 2a-b, Supplementary Fig. 4b**). We then examined the three canonical CRC driver genes (*APC, KRAS* and *TP53*, hereafter abbreviated as AKP) and defined AKP and non-AKP crypts as those with or without AKP somatic alterations, respectively (**Fig. 2b)**.

**Figure 2.**
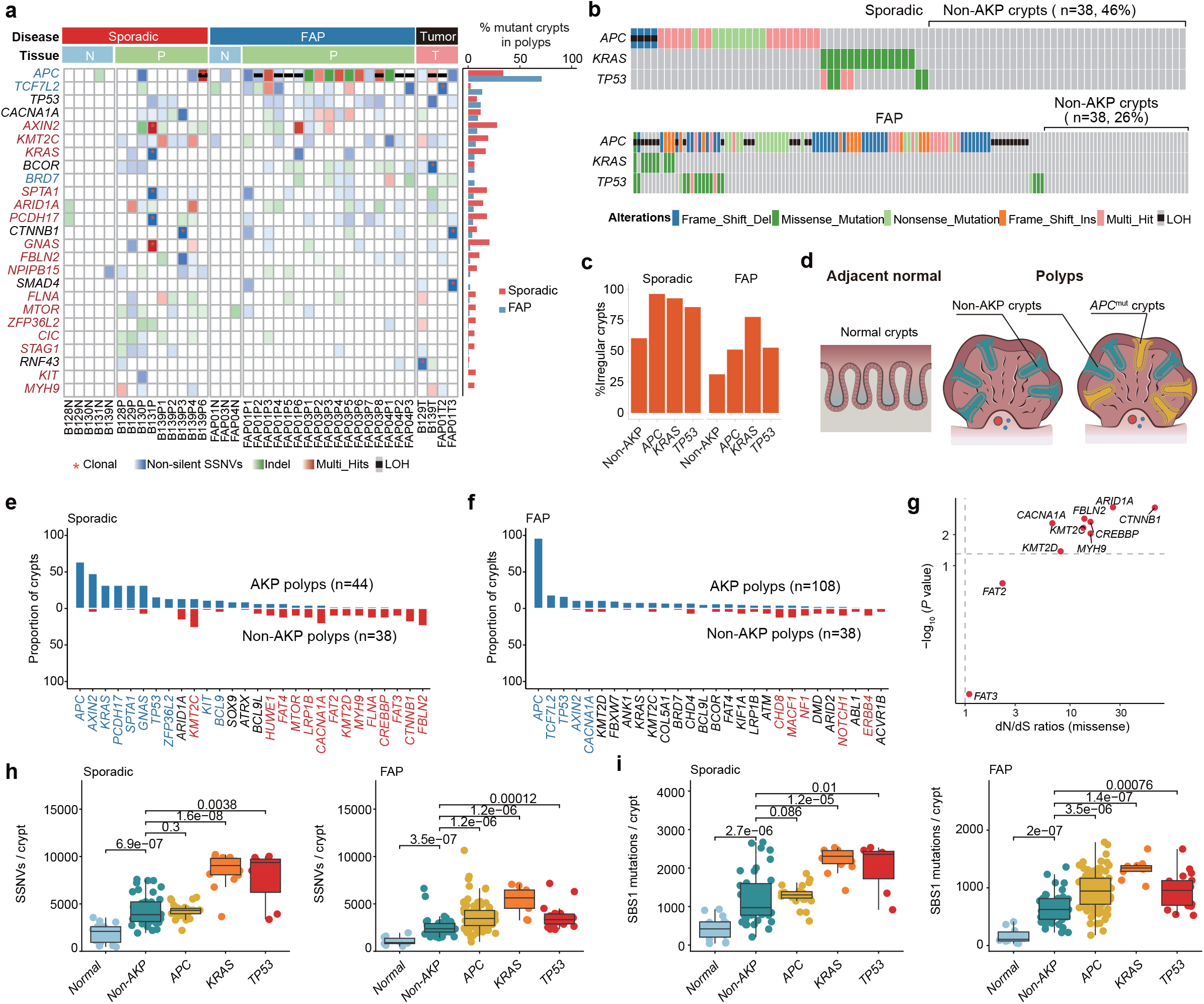
Distinct driver gene landscape of sporadic and hereditary colorectal polyps. **a**, Non-silent somatic single nucleotide variants (SSNVs), insertions/deletions (indels) and loss of heterozygosity (LOH) in putative driver genes. The color gradient of each box denotes the mutational clonality in the corresponding tissue or normal tissue. A mutation was defined as clonal if present in more than 90% of crypts within a tissue containing at least five single-crypts with WGS data. **b**, Oncoplot summarizing the landscape of non-silent SSNVs, indels and LOH within canonical APC/KRAS/TP53 (or AKP) genes across single polypous crypts from sporadic (the top) and FAP (the bottom) patients. **c**, The proportion of irregular crypts in each indicated group from sporadic (left) and FAP (right) patients. **d**, Schematic illustrating that each premalignant polyp consists of mixed non-AKP and AKP crypts. **e-f**, Prevalence of driver gene mutations in non-AKP versus AKP crypts from sporadic (**e**) and FAP (**f**) patients. *P* value, Fisher’s exact test. Genes highlighted showing *P* < 0.05, or fold change > 1.5 with at least 10% in non-AKP or AKP crypts. **g**, dN/dS ratios of missense mutations in the significantly enriched genes in sporadic polypous crypts (**e**). *P* value, log likelihood-ratio test. Vertical and horizontal lines correspond to absolute dN/dS values of 1 and P < 0.05, respectively. **h-i**, Comparison of the burden of SSNVs (**h**), SBS1-associated mutations (**i**) in normal, non-AKP and AKP polypous crypts from sporadic (left) and FAP (right). *P* value, two-sided Wilcoxon rank-sum test.

Strikingly, nearly half (46% or 38 out of 82) of the sporadic premalignant crypts were non-AKP, whereas this proportion was much lower in FAP (26%, 38 out of 146, **Fig. 2b**). Importantly, non-AKP crypts in sporadic polyps often showed more irregular morphology than those in FAP polyps (**Fig. 2c-d**), indicating that these non-AKP crypts were bona fide dysplastic crypts. In sporadic polyps, putative driver genes including *KMT2C, CACNA1A, FAT2, CTNNB1, FBLN2, etc* were more frequently mutated in non-AKP crypts than in AKP ones (**Fig. 2e**). However, few genes were found in FAP polyps for this comparison (**Fig. 2f**). In fact, these driver genes like *KMT2C, CACNA1A* and *FBLN2* also showed stronger selection in sporadic polyps compared to normal mucosal crypts (**Fig. 2g, Supplementary Fig. 4c-d**), suggesting they confer a fitness advantage during the earliest stages of initiation. These findings indicate that early sporadic adenomas are often initiated through non-AKP CRC driver mutations, whereas adenomas in FAP patients tend to follow the *APC* double-hit trajectory because of the germline heterozygous *APC* background.

We then examined whether somatic drivers in sporadic and FAP polyps were associated genomic evolutionary trajectories by subgrouping premalignant crypts into non-AKP, *APC*-mutant, *KRAS*-mutant or *TP53*-mutant ones (**Fig. 2h-i, Supplementary Fig. 5**). Interestingly, we found that non-AKP polypous crypts harbored a comparable number of somatic mutations with *APC*-mutant crypts but fewer than *KRAS*-mutant or *TP53*-mutant crypts in sporadic patients (**Fig. 2h**). This pattern was also reflected in clock-like mutations (SBS1 and SBS5) (**Fig. 2i, Supplementary Fig. 5b**). These data suggest that, in sporadic tissues, non-AKP and *APC*-mutant clones may expand in parallel during early lesion development. In FAP, by contrast, *APC*-inactivated crypts likely represent a more advanced stage than non-AKP crypts. Together, these results further show that sporadic and hereditary precancers have deterministic yet distinct trajectories towards CRC.

### Somatic *APC* mutations often arise prenatally in FAP patients

To determine when key driver mutations arise, we estimated the timing of somatic events along the phylogenetic trees using a molecular clock model, as in a previous study^48^. We reconstructed single-crypt WGS phylogenies at the patient level, including all crypts from different polyps of the same patient. Somatic mutations were assigned to corresponding branches and timing of the events was inferred as the age from fertilization, while accounting for patient-specific and clade-specific mutation rates and the elevated mutation rate during embryonic development^49^ (**Fig. 3, Supplementary Table 4, Methods**). Since somatic mutations accumulate approximately linearly with age along each branch, mutations located on upper branches of a phylogenetic tree indicate their earlier acquisition (**Methods**).

**Figure 3.**
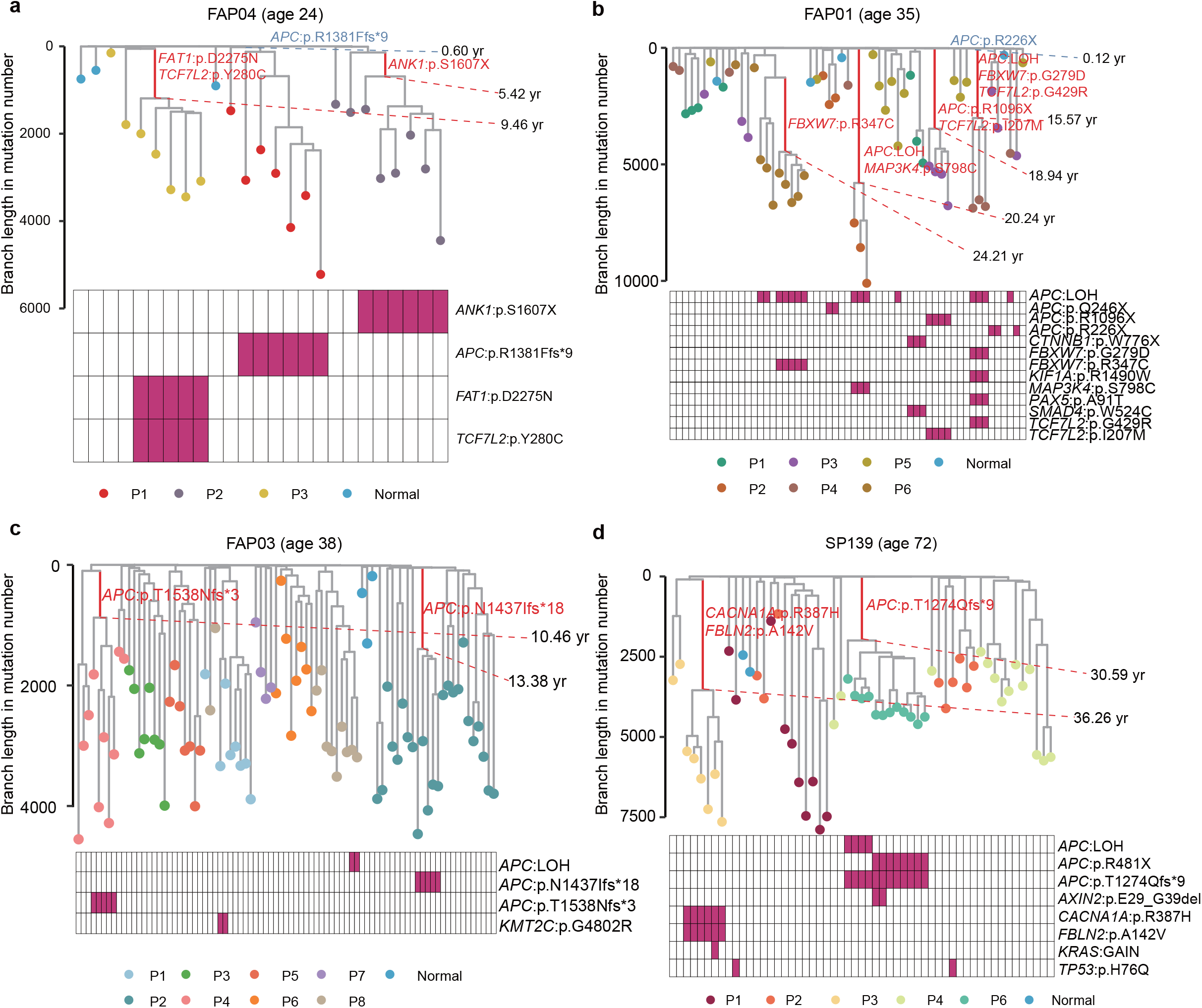
The timing of driver mutations in premaligant polyps from 4 patients. **a-d**, Time-resolved phylogenetic tree of individual crypts from three FAP (**a-c**) and one sporadic patient (**d**). Somatic driver mutations and the timing of occurrences are highlighted along corresponding branches. The tips of the branches represent individual crypts. Mutations that likely drive clonal expansion are labeled in red. Inferred patient ages (yr, years after conception) of the most recent common ancestor (MRCA) harboring driver mutations are depicted.

Strikingly, two out of three FAP patients (FAP04 and FAP01) acquired the first somatic *APC* mutation within the first year of life from fertilization. Specifically, FAP04 acquired the *APC*^*R1381Ffs*9*^ mutation at 0.6 years (7.2 months) old from zygote (95% confidence interval or CI: 0.45–0.87) (**Fig. 3a**), and FAP01 acquired the *APC*^*R226X*^ mutation at 0.12 years (1.4 months) old from fertilization (95% CI: 0.08–0.16) (**Fig. 3b**). In FAP01, the somatic *APC*^*R1096X*^ mutation and two *APC* 5qLOH events occurred at about 15.57 (95% CI: 15.12–16.09), 20.24 (95% CI: 19.90– 20.58), and 18.94 (95% CI: 18.41–19.68) years old, respectively (**Fig. 3b**). In FAP03, two somatic *APC* mutations occurred at about 10.46 (95%: 8.77–14.57) and 13.38 (95% CI: 12.27– 16.22) years old, respectively (**Fig. 3c**). In the sporadic case (SP139, 72 years old), the acquisition of *APC*^*T1274Qfs*9*^ occurred at about 30.59 (95% CI: 23.28–38.38) years old (**Fig. 3d**), over 40 years prior to diagnosis. Somatic *APC* was indeed associated with clonal expansions according to the single-crypt phylogenetic trees. However, we also observed non-AKP driver-associated clonal expansions, including *CACNA1A* and *FBLN2* in SP139, and *FAT1* and *TCF7L2* in FAP04.

### Non-AKP clones contribute to polyclonal origins in sporadic polyps

The polyclonal origin of human colorectal premalignant tissues has recently been demonstrated as a new paradigm^18,19^. We therefore asked how AKP and non-AKP clones contribute to polyclonality in sporadic and FAP polyps. To this end, we reconstructed single-crypt phylogenies for each tissue, along with matched normal crypts from the same patient (**Fig. 4a-b, Supplementary Fig. 6**). A clonality score was defined based on polygenetic topology to quantify the clonality and distinguish polyclonal (score <0.05) from monoclonal (score >=0.05) polyps (**Methods**).

**Figure 4.**
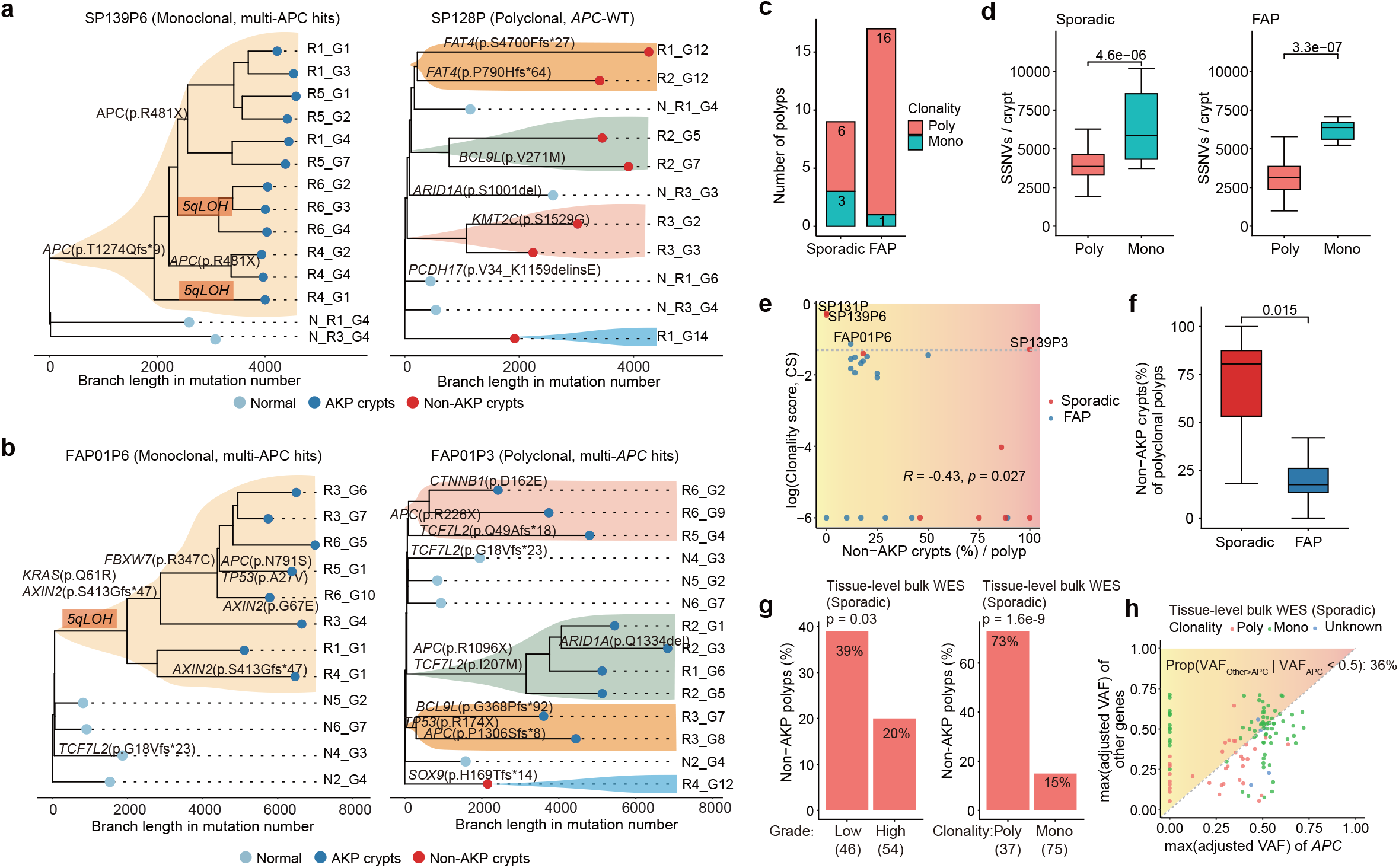
Non-canonical driver clones fuel the polyclonal origins of sporadic polyps. **a-b**, Single-crypt phylogeny for representative monoclonal (left) and polyclonal (right) polyps from sporadic (**a**) and FAP (**b**) patients. **c**, Number of polyps exhibiting polyclonal and monoclonal origins from sporadic and FAP patients. **d**, The burden of SSNVs per crypt in polyclonal and monoclonal polyps from sporadic (left) and FAP (right) patients. **e**, Correlation between the fraction of non-AKP crypts and clonality score for each polyp. **f**, Fraction of non-AKP crypts in polyclonal polyps from sporadic (left) and FAP (right) patients. **g**, Non-AKP polyps are enriched in low-grade and polyclonal tissues based on tissue-level bulk WES data of 107 published premalignant sporadic polyps^19^ and 10 newly collected ones. **h**, Adjusted variant allele frequency (VAF) of *APC* and other mutated genes in each sporadic polyp. Pearson’s *r* and *P* value are shown in (**e**) and (**h**). *P* value, two-sided Wilcoxon rank-sum test (**d**,**f**) or Fisher’s exact test (**g**).

Indeed, most premalignant polyps (85% or 22 out of 26) were composed of multiple clonal lineages and had a low clonality score (**Fig. 4a-b, Supplementary Figs. 6-7**), supporting their polyclonal origins. Interestingly, polyclonal origin was more common in FAP than in sporadic polyps (**Fig. 4c**). This is likely due to sporadic polyps being diagnosed later than FAP, as reflected by the more irregular morphology of single crypts in sporadic polyps compared to FAP polyps (**Fig. 2d-e**). Polyclonal tissues harbored fewer somatic mutations than monoclonal ones, consistent with progressive polyclonal-to-monoclonal evolution^19^ (**Fig. 4d**). The fraction of non-AKP crypts within a tissue negatively correlated with the clonality score (**Fig. 4e**), indicating that non-AKP clones were strongly enriched in polyclonal tissues. This pattern was more prominent in sporadic polyclonal tissues, with a substantial fraction of non-AKP crypts (10–100%; **Fig. 4f**). In contrast, FAP polyclonal tissues were often characterized by parallel evolution of distinct lineages carrying multiple independent somatic *APC* alterations (31% or 5 out of 16 tissues, **Fig. 4b, Supplementary Fig. 6**).

To validate the contribution of non-AKP clones in sporadic polyps, we analyzed a large tissue-level bulk WES cohort comprising 107 sporadic premalignant lesions^19^ and 24 newly collected lesions in this study (**Supplementary Tables 1 and 5**). Here polyclonal status was identified based on a skewed distribution of variant allele frequencies (VAFs) with a mean VAF significantly lower than 0.5. Non-AKP tissues more often had low-grade and polyclonal origins (**Fig. 4g**), and a substantial subset of polyps (36%) were dominated by non-*APC* clones (**Fig. 4h**).

Together, these results demonstrate that the polyclonal origins of sporadic and hereditary polyps are genetically different. While parallel acquisition of somatic *APC* alterations contributes to polyclonality in FAP polyps, sporadic polyps consist of multiple non-AKP clones.

### Monoclonal transition recapitulates Big-Bang-like growth in colorectal precancer

Since the spatial information where each crypt was dissected from the tissue was recorded, we next performed a phylogeographic analysis to distinguish spatial lineage segregation from lineage intermixing. Spatial segregation refers to the scenario where phylogenetically close crypts are located in neighboring regions, as exemplified by sporadic (SP130P) and FAP (FAP03P4) cases (**Fig. 5a, Supplementary Figs. 8-9**). In contrast, spatial lineage intermixing refers to the scenario where phylogenetically close crypts are found in spatially distant regions, as exemplified by a sporadic case (SP131P) and an FAP (FAP01P6) case (**Fig. 5b, Supplementary Figs. 8-9**). In these tissues, the presence of the same low-VAF somatic mutations across distant regions further supported spatial lineage intermixing (**Fig. 5c, Supplementary Fig. 9c-d**).

**Figure 5.**
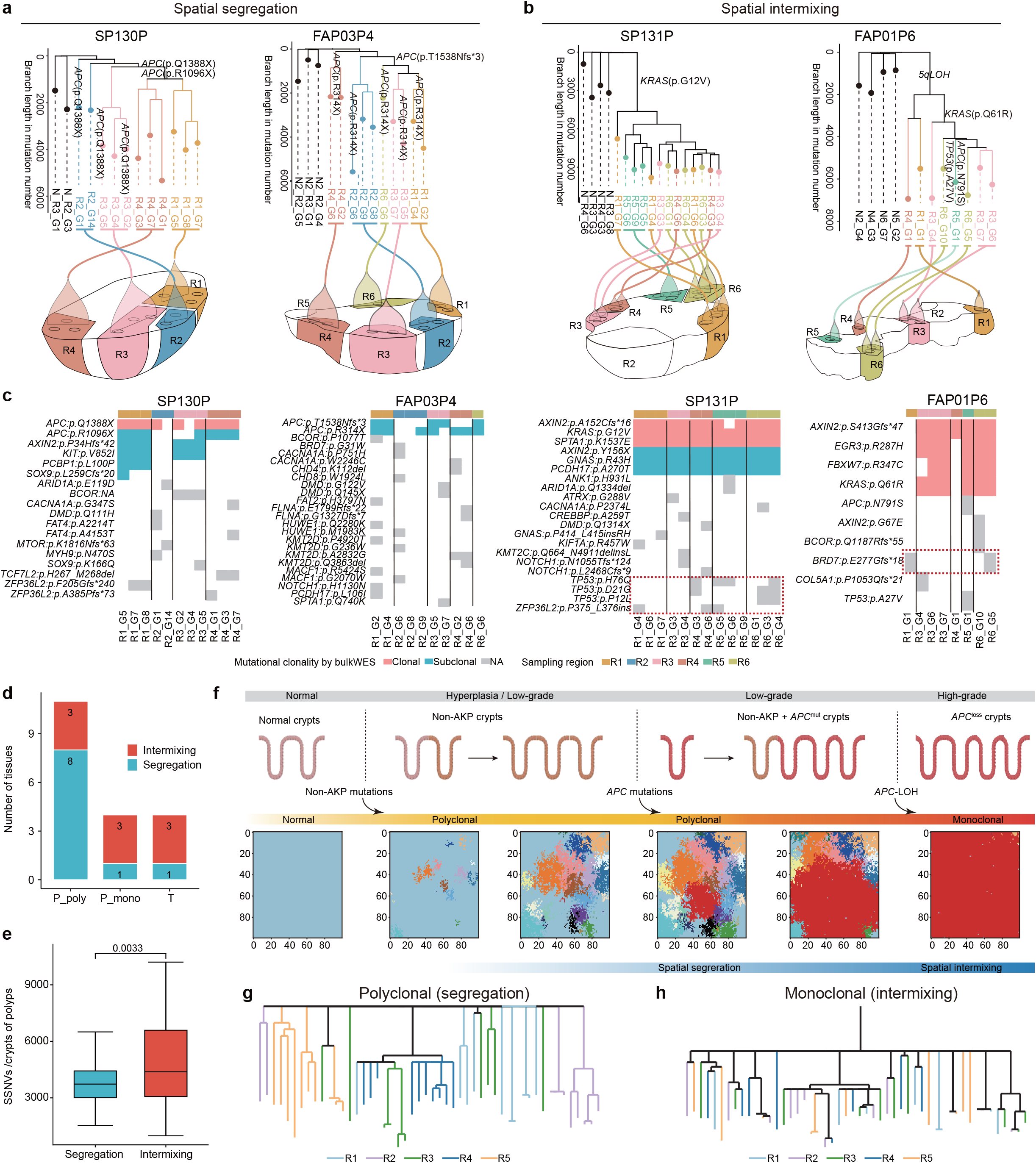
Phylogeographic analysis reveals the onset of Big-Bang growth in colorectal polyps. **a-b**, Single-crypt phylogeny (top) with corresponding spatial locations annotated (bottom) for representative polyps exhibiting spatial lineage segregation (**a**) or lineage intermixing (**b**). **c**, Heatmaps indicating the presence of representative clonal and subclonal mutations in CRC-associated driver genes across individual crypts per polyp. Mutational clonality (clonal or subclonal) is determined by tissue-level bulk WES (Methods). NA, present in single-crypt WGS data but non-detected in tissue-level bulk WES data. **d**, The number of tissues with spatial lineage intermixing or segregation in polyclonal polyps (P_poly), monoclonal polyps (P_mono) and CRCs (T). **e**, The burden of SSNVs per crypt in polyps with spatial lineage segregation versus lineage mixing. *P* value, two-sided Wilcoxon rank-sum test. **f**, A 2D simulation model of early tumor growth following polyclonal-to-monoclonal transiton. Initially, multiple clones carrying distinct non-AKP mutations (different colors) emerge from normal tissue, leading to a polyclonally growing tumor. Then a clone harboring a dominant driver mutation (e.g. *APC* mutation, red clone), confers a higher selective advantage leading to outgrowth and monoclonal transition. **g-h**, Phylogenetic trees reconstructed from the mutation profiles in simulated crypts sampled from the predefined spatial regions during the polyclonal (**g**) and monoclonal (**h**) states. Different colors indicate distinct sampling regions.

Interestingly, we found that the pattern of spatial lineage segregation versus intermixing was strongly associated with the tissue clonality (polyclonal versus monoclonal) (**Fig. 5d**). Specifically, spatial lineage intermixing was more prevalent in monoclonal polyps (75% or 3 out of 4) than in polyclonal polyps (27.3% or 3 out of 11) (**Fig. 5d**). Polyps with spatial lineage intermixing carried significantly more somatic mutations at the single-crypt level than polyps with spatial lineage segregation (**Fig. 5e**), indicating that intermixing marks a more advanced evolutionary stage. Notably, most malignant CRCs (75%, 3 of 4) in our cohort also exhibited spatial lineage intermixing (**Fig. 5d** and **Supplementary Fig. 10**). Since spatial lineage intermixing is a defining feature of Big-Bang model of tumor growth^31^, our data indicate that Big-Bang-like growth can begin as early as the premalignant stage.

To further verify spatial lineage intermixing as a key feature of Big-Bang tumor growth, we simulated the two-dimensional spatial growth of virtual tumors. Then, we sampled individual crypts from multiple regions to reconstruct single-crypt phylogenetic trees, mimicking our multi-regional single-crypt sampling strategy (**Methods**). Our simulations revealed strong spatial segregation for the early initiated clones through polyclonal origins, followed by a selective sweep of a dominant clone (**Fig. 5f**). Importantly, this clonal sweep generates the characteristic spatial intermixing pattern as observed in the monoclonal premalignant tissues and CRCs (**Fig. 5f-h** and **Supplementary Fig. 11**). Together, our phylogeographic analysis and simulations revealed that spatial lineage architecture is mechanistically linked to the disease stage, and that the polyclonal-to-monoclonal transition in precancerous tissues aligns with Big-Bang-like tumor growth.

## Discussion

Although both sporadic and hereditary colorectal precancers often arise through polyclonal origins, our single-crypt WGS analysis shows that they initiate and evolve through distinct genomic routes. Sporadic polyps frequently develop not by the canonical driver genes (e.g. *APC, KRAS* and *TP53*), but instead through non-AKP driver mutations including *KMT2C, CACNA1A and FBLN2*. In contrast, the second hit on the remaining allele of *APC* in FAP lesions contributes to polyclonal initiation, where most premalignant crypts hit by a second *APC* mutation or 5q LOH at a strikingly early in life (often prior to birth). These support the two distinct molecular trajectories underlying the same mode of precancer polyclonality observed in both sporadic and hereditary conditions.

Canonical CRC drivers, including *APC, KRAS*, and *TP53*, are thought to be essential for tumorigenesis; however, one important implication of our findings is that early polyclonal expansion in polyps not necessarily require those CRC drivers. In sporadic lesions, non-AKP crypts also showed dysplastic morphology and high mutation burdens comparable to those of *APC*-mutant crypts. This suggests that such clones are not passive bystanders, but active participant in lesion clonal expansion. One plausible model is that these mutations act as “mini-drivers”: individually modest, but sufficient to disrupt epithelial homeostasis, alter cell competition, or reshape local microenvironmental interactions^6,19,50,51^. Their functions might require specific timing, such as the short time window before monoclonal sweep. Our data therefore expand the current model of colorectal tumor initiation from a strictly gene-centric sequence to a broader ecological and evolutionary process.

Tumor growth and progression involve the breakdown of normal tissue organization^52^. Previous studies showed the Big Bang model and exposes variegation in carcinomas but not adenomas^31^. However, we found Big-Bang growth occurs early at monoclonal transition of polyps, exhibiting a spatially mixed architecture and substantial variegated branch mutations. Notably, early polyclonal polyps display a spatially segregated pattern, occasionally accompanied by local monophyletic clusters, suggesting selective advantages in clonal competition may be the basis under field cancerization. These observations recapitulate predictions of a Big-Bang growth and directly support the concept that tumors are “born-to-be-bad”. This underscore that tumor-initiating events in colorectal cancer may occur much earlier than previously recognized, which may help to better inform strategies for early detection, risk stratification, and prevention before these morphologically normal crypts transform into neoplasia. Collectively, our study defines distinct genomic trajectories of early colorectal tumorigenesis in sporadic and hereditary contexts, providing a conceptual framework for understanding how these lesions diverge during initiation and converge when out. Our findings support a view in which precancer initiation is shaped by both somatic mutation and tissue organization. The former is reflected by the distinct driver landscapes of sporadic and FAP lesions, whereas the latter is captured by the spatial restructuring that accompanies the transition from polyclonal to monoclonal stage.

These new insights may also inform clinical practice in early cancer interception. As monoclonal polyps with Big-Bang-like lineage mixing may mark a more advanced and potentially higher-risk stage of disease, defining the biomarkers of this stage would greatly help guide early intervention. Colorectal cancer also develops through other routes, including Lynch syndrome and colitis-associated neoplasia in inflammatory bowel disease^11^. Defining the evolutionary trajectories of these additional pathways will complete our understanding of CRC initiation across multiple disease subtypes and guide earlier detection and prevention.

## Methods

### Sample collection

Our human colorectal precancer cohort consisted of nine patients (six sporadic and three with FAP) (**Supplementary Table 1**), recruited from the Sixth Affiliated Hospital of Sun Yat-sen University. All of the sporadic patients had synchronous polyps and CRC located at distinct sites of the colon. Of the three patients with FAP, two had developed CRC by the time tissue samples were collected. All biospecimen collection protocols were conducted in accordance with the principles of the Declaration of Helsinki and were approved by the Institutional Review Board (IRB) of the Sixth Affiliated Hospital of Sun Yat-sen University (2019ZSLYEC-06). All enrolled patients had signed informed consent forms, and all samples were anonymously coded in accordance with local ethical guidelines. In total, we have obtained 28 polyps (P), 9 synchronous colorectal tumors (T), and 17 adjacent normal tissues (N) from these 9 patients (**Supplementary Table 1**). Of these 54 samples, 46 samples, including normal mucosa (n = 16), premalignant polyps (n = 26), and synchronous malignant adenocarcinomas (n = 4), were subject to single-crypt isolation and WGS, while the remaining 8 samples were subject to tissue-level bulk WES (**Supplementary Table 1**).

The bulk tissue samples were immediately frozen and then subjected to histopathological assessment, single-crypt dissection, WGS, and/or tissue-level bulk WES. For the histopathological assessment, the sample tissues were embedded in an Optimal Cutting Temperature (OCT) compound (Sakura Finetek, USA) using pre-cooled molds. Then, cryosectioning, fixation, and hematoxylin and eosin (H&E) staining were conducted.

### Single-crypt isolation and morphology assessment

We adapted the single-crypt isolation procedure from the published protocols^19,30^ (**Fig. 1b**). First, four to eight spatially distinct regions were dissected from each tissue (**Supplementary Fig. 1**). Then, the dissected tissues were placed on a clean glass slide under the microscope and carefully added 20 μL pre-cold PBS. Next, individual crypts (or glands) were manually dissociated using a 23G (0.6 x 25 mm) injection needle and aspirated using a pipette. Each crypt was then transferred to a PCR tube containing 20 μL of proteinase K buffer.

Based on microscopic morphology, crypts obtained through microdissection can be classified as irregular crypts^53^ if they meet at least one of the following criteria: 1) crypts exhibiting significant enlargement, including elongation and increased diameter. Specifically, the length of an irregular crypt is approximately twice that of adjacent normal crypts^54^, and the diameter of irregular crypts in aberrant crypt foci (ACF) measure is up to 1.5 times greater than that of a normal crypt^55^; 2) Crypt basal dilatation ≥ 10% of crypts^56^; 3) Irregular crypts displaying double or triple branching crypts, or other pronounced forms of budding and branching^54-56^; 4) A peculiar growth pattern involving the horizontal orientation of deep crypts that seem to grow parallel to the muscularis mucosae, often creating an inverted T- or L-shaped crypt^56^.

### Single-crypt whole genome sequencing

Genomic DNA was extracted from isolated crypts by following a published protocol^33^. First, each crypt was transferred to a pre-loaded 200 μL PCR tube containing 10 μL of lysis buffer (30 μL Tris-HCl (pH 8.0), 5 μL Tween-20, 5 μL IGEPAL CA-630, 50 μL 25 ug/mL proteinase K, and 910 μL ddH_2_O). The tubes were then incubated at 50 °C for 12 minutes. The DNA was captured using 100 μL of a 1:1 mixture of AMPure beads (Beckman Coulter, USA) and TE buffer (pH 8.0). The beads were washed twice with 150 μL of 80% (vol/vol) ethanol for one minute. Finally, the genomic DNA was eluted in 12 μL of ddH_2_O and quantified using a Qubit 4.0 Fluorometer (Invitrogen, US).

Since a classical crypt contains approximately 2000 cells^57^, we selected crypts with 1∼20 ng of total DNA to avoid unexpected situations, such as aspiration of other tissues (resulting in excessive total DNA) or failure to aspirate the crypts (resulting in insufficient total DNA). The low-input DNA sequencing protocol was adapted from a previously published method^33^. Based on the measured DNA concentration, we prepared libraries using either the Vazyme TruePrep DNA Library Prep Kit V2 (TD501/502/503, Vazyme, China) or the Hieff NGS® OnePot Pro DNA Library Prep Kit V2 (Yeason, China), according to the manufacturer’s instructions. Each library was assigned a unique index pair and was further validated by a TOPO cloning followed by Sanger sequencing. Library quality was check by TapeStation (Agilent) or qSep (BiOptic). Qualified libraries were sequenced using the NovaSeq 6000 (Illumina, USA) or Novaseq X plus platforms (Illumina, USA). Each sample has at least 100 GB of raw sequencing data to ensure further analysis.

### Tissue-level bulk WGS and WES

Tissue-level bulk WGS was performed on eight normal samples to serve as controls for single-crypt WGS. WES was performed on nine normal samples and 33 polyps and tumor tissues to validate somatic mutations. First, genomic DNA was extracted from fresh-frozen samples using the DNeasy® Blood & Tissue Kit (Qiagen, USA) and quantified it using the Qubit® 4.0 Fluorometer (Invitrogen, USA). We constructed DNA libraries for samples with a DNA yield of >200 ng. WGS libraries were prepared using the Rapid Plus DNA Library Prep Kit for Illumina (RK20208), following the manufacturer’s guidelines. The libraries were then sequenced on the Illumina NovaSeq 6000 or NovaSeq X Plus to a depth of 28x. WES libraries were prepared using the NEBNext® Ultra™ DNA Library Prep Kit (NEB) following the manufacturer’s recommendations. Index codes were added, and the libraries were captured using the SureSelect XT Human All Exon V6 kit (Agilent Technologies). The libraries were sequenced on the Illumina NovaSeq X Plus to a depth of 190x.

### Alignment for WGS and WES data

The genomic sequencing data were preprocessed using an in-house pipeline, as previously described^19,34,35^. In brief, FASTQ files were processed using fastp v0.12.4 (-q 5 -u 50 -w 2 -l 50 -n 5 -z 6) and mapped to the human reference genome (GRCh38) using BWA v0.7.17-r1188 (bwa mem -M -t 7 -R). Following GATK best practices and the associated set of tools v4.2.0.0^58^, duplicated reads were marked in sorted bam files using MarkDuplicates (VALIDATION_STRINGENCY=LENIENT), called bases were recalibrated using BaseRecalibrator against the indel references (Mills_and_1000G_gold_standard.indels.hg38.vcf.gz), the SNPs from dbSNP146 (dbsnp_146.hg38.vcf.gz) and 1000 Genomes Project (1000G_phase1.snps.high_confidence.hg38.vcf.gz). Finally, the final bam files were generated using ApplyBQSR. Sequencing depth was calculated using mosdepth v0.3.3 for WGS and bamdst v1.0.9 (https://github.com/shiquan/bamdst) for WES.

### Somatic mutation calling and filtering

As previously described^19,34,35^, Mutect2 was used to identify somatic mutations in tumor or polyp crypts with the default options, using the matched tissue-level bulk adjacent normal tissue as germline control for each individual. We filtered out artifactual and background germline variants using the gnomAD and customized PoN germline resources. Raw somatic variants were filtered using GATK’s FilterMutectCalls with the default settings, and variants labeled “PASS” were retained for further analysis. Additionally, somatic mutations were called separately on single-crypt WGS and tissue-level bulk WGS data using Strelka v2.9.2, and candidate indels were called with Manta v1.6.0. To eliminate false-positive mutation calls from the single-crypt WGS data, we retained only variants called by both Mutect2 and Strelka v2.9.2. We obtained a consensus variant call set using VariantFilter (https://github.com/rschenck/VariantFilter). Finally, high-confidence mutations were defined as having a VAF greater than 0.15, at least two supporting variant reads, at least five total reads in crypt samples, and no supporting variant reads in the matched tissue-level bulk normal sample.

Moreover, we specifically recalled a subset of mutations in 367 putative CRC driver genes, which integrate our prior assembled gene list for Pan-cancer and/or CRC cohorts^35^, newly CRC-specific drivers annotated by Chinese cohort^59^ and additional precancer-specific drivers^5^. To avoid potential artifacts in the single-crypt WGS data, a blacklist of abnormal samples was curated and excluded from the downstream analysis by following criteria: 1) an abnormally high C>A mutation ratio, 2) unusual signature distribution, and 3) multiple hits within numerous drivers. Additionally, 20 polypous crypts with mutation burdens comparable to normal crypts from the same patient were classified as “normal-like” (**Supplementary Table 3**) and removed from downstream analyses.

Somatic mutations from the matched tissue-level bulk WES data were detected using our previously established pipeline^19,34,35^. Mutations at tissue-level with an upper bound of the 95% CCF confidence interval ≥ 1 were classified as ‘clonal’, and others as ‘subclonal’.

### Somatic copy number alterations and LOH analysis

SCNA, ploidy, and tumor purity for single-crypt or tissue-level bulk lesions were estimated using TitanCNA^60^, a hidden Markov model-based method. For each patient, we identified germline heterozygous SNP at dbSNP 146 loci in the normal biopsy using SAMtools and SnpEff. Read counts for 10,000 base pair (bp) bins across the genome were generated for all samples using HMMcopy. Then, TitanCNA was used to determine the allelic ratios at germline heterozygous SNP loci in normal, precancerous and cancerous crypts and the depth ratios between crypts and control normal samples in the bins containing those SNP loci. Nine TitanCNA runs with nine combinations of subclone number, purity, and ploidy were performed. Different numbers of subclones (n = 1, 2 or 3), purities (n = 0.2, 0.4, or 0.6), and ploidy (2) were set. One run was selected for each tumor biopsy based on manual inspection of the fitted results. Results with a single subclone were prioritized unless results with multiple subclones better fit the data visually. Notably, assuming the crypt as a single clone, we also re-adjusted the output using an in-house script. SCNAs were identified in single-crypts relative to paired normal tissue-level bulk tissues. For amplification, the absolute copy number was greater than 2.8, and the copy number relative to median ploidy was greater than 0.8. For deletions, the absolute copy number was less than 1.2 and copy number relative to median ploidy was less than -0.8. The neutral LOH regions were called using TitanCNA. These putative CNAs were also detected using the Sequenza v3.0.0 package for R, as previously described^19^.

### Germline *APC* mutation calling

To identify *APC* germline mutations in the three FAP patients, we first extracted reads from the *APC* region (chr5:112,737,885-112,846,239) using samtools v1.12 and called variants with bcftools v1.14. Then, we merged and annotated the mutations identified across all samples from each patient. An *APC* mutation was considered a germline mutation if it met the following criteria: (1) non-silent; (2) present in the 1000 Genomes or ExAC database with a minor allele frequency (MAF) of less than 1%; (3) recurrent in nearly normal crypts and the matched tissue-level bulk samples with a variant allele frequency of greater than 20%; and (4) detected in approximately 90% of preneoplastic or neoplastic crypts.

### Mutational signature extraction and assignment

Mutational signatures were identified using previously reported piplines^61^. First, a hierarchical Dirichlet process (HDP v0.1.5)^37^ was used without priors to extract the mutational signatures. We set the alpha and beta to one for the alpha clustering parameter. Gibbs sampler was ran with 20,000 burn-in iterations (parameter ‘burnin’). With a spacing of 1000 iterations (parameter ‘space’), 100 iterations were collected (parameter ‘n’). After each Gibbs sampling iteration, three iterations of concentration parameter sampling (parameter ‘cpiter’) were performed.

For identified mutational signatures, only the signatures with ≥0.9 cosine similarity with the reference as the same signatures were considered. For the remaining signatures, HDP was ran with all known PCAWG reference signatures V3.1^62^ as priors and kept the extracted signatures as a shortlist of candidate reference signatures. To deconvolute composite signatures and to equate obtained HDP signatures to reference signatures, we used an expectation-maximization algorithm to deconstruct these signatures into reference constituents. We ran a second round of expectation maximization that only kept reference signatures with >10% contributions for each HDP signature to reduce overfitting. The final SBS mutational signatures permitted in each patient were the corresponding deconvoluted reference signatures for HDP components contributing to at least 5% of mutations in at least one sample. This amounted to reference signatures SBS1, SBS2, SBS5, SBS13, SBS18, SBS88 for normal crypt and SBS1, SBS2, SBS3, SBS5, SBS13, SBS18, SBS88, SBS3, SBS17a, SBS17b for polyp and tumor crypts. The final proportion of reference signatures was assigned to each crypt using sigfit (v2.2)^38^.

### Somatic LINE-1 retrotransposition analysis

The LINE1 (L1) retrotranspositions was called using DELLY^46^ and xTea^45^ with matched tissue-level bulk normal samples and single crypts to detect somatic insertions. Potential germline calls, overlapping with events found in matched tissue-level bulk samples, were removed.

For DELLY, we excluded rearrangements with poor mapping quality (median MAPQ < 40), insufficient supporting reads, or excess discordant reads in matched normal BAM files. We also removed short deletions and duplications (<1 kb) lacking soft-clipped reads and unbalanced inversions <5 kb with fewer than five supporting reads, as these are typically library artifacts. Breakpoint positions and microhomology sequences were refined using SA tags from soft-clipped reads. Breakpoint variant allele fraction was estimated from discordant read pairs supporting the rearrangement relative to concordant read pairs supporting the wild-type configuration.

For xTea, insertion-calling thresholds were adjusted according to insertion clonality and sequencing depth. Candidate insertions with few or no support in the matched normal sample were classified as somatic. Insertions overlapping repetitive regions from the same TE family were removed when the reference repeat showed low divergence from the RepeatMasker consensus. xTea then required consistent clustering of 5′ and 3′ clipped reads and their corresponding discordant mates within the expected insert-size range. Candidates supported on only one side of the breakpoint were further resolved using discordant reads or local read-depth patterns to identify transductions or target-site deletions.

### dN/dS analysis

We used the R package dndscv^63^ (https://github.com/im3sanger/dndscv) to look for evidence of positive selection in our dataset. The dndscv package compares the observed ratio of missense, truncating and nonsense to synonymous mutations with that expected under a neutral model. It incorporates information on the background mutation rate of each gene and uses trinucleotide-context substitution matrices. The approach provides a global estimate of selection in the coding variant dataset, from which the number of excess protein coding, or driver mutations can be estimated. In addition, it identifies specific genes that are under significant positive selection.

### Assignment of mutations to branches

We used Sequoia^64^ to reconstruct the phylogenetic trees and assign mutations to branches, which were then used in the timing analysis. First, Sequoia uses a maximum parsimony framework implemented in MPBoot with default settings to infer the topology of the phylogenetic tree. Then, Sequoia uses an expectation maximization method in the treemut^48^ package to assign mutations to the phylogenetic trees and estimate branch length.

The treemut is based on Bayes rule: *P*(*mutation has genotype*) = *P*(*read depth*|*genotype*) ⋅ *P*(*genotype*). Each mutation has a corresponding true genotype vector, in which the elements are 1 for crypts descending from a particular ancestral branch and 0 for those that do not. The prior probability of a mutation with a genotype *P*(*genotype*) is proportional to the branch length. Read depths are modeled using a binomial distribution with sample-specific error rates and an assumed variant allele frequency *VAF* = 0 for wild-type sites and *VAF* = 1/*ploidy* for variant sites. This process is iterated, updating the branch lengths with each iteration. After estimating the maximum likelihood probability that each mutation belongs to each branch, treemut hard-assigns mutations.

### Timing of somatic driver mutations on phylogenetic trees

Based on the phylogenetic trees reconstructed by Sequoia, we used rtreefit to date the timing of somatic driver mutations. rtreefit converts branch lengths in molecular time, number of mutations, to branch length in units of time (years) and jointly fitted wild type rates, mutant rates and absolute time branch lengths using a Bayesian per patient tree-based model under the assumption that the observed branch lengths are Poisson distributed with *mean* = *branch duration* × *sensitivity* × *mutation rate* and subject to the constraint that the root to tip duration is equal to the age at sampling. Furthermore, the method accounts for an elevated mutation rate during embryogenesis by assuming an excess mutation rate through development. Models were fitted across four chains each with 20,000 iterations. The trees are guaranteed to have a root to tip distance that matches the sampling age of the colony.

In detail, rtreefit considers a rooted tree in which each edge *i* consists of an observed mutation count, *m*_*i*_ and a true duration, *t*_*i*_. A given edge and its child node are denoted by the same label. *D*(*i*) is the tips descending from node *i* and *A*(*i*) is the ancestral node excluding the root. If each tip of the tree has a known time *T*_*k*_, then we have the following constraint applies:

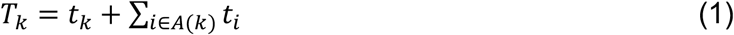

and *T*_*k*_> *t*_*i*_ > 0.

rtreefit incorporates this constraint by optimizing the interior branches of the tree with reparametrized branch lengths *x*_*i*_ transformed to be in the range 0 < *x*_*i*_ < 1, where *x*_*i*_ can be thought of as stick breaking fractions. If *j* is an edge whose parent node is the root then:*t*_*i*_ = *x*_*i*_(*T*_*k*_: *k* ∈ *D*(*j*)).

For other interior edges *i* we have

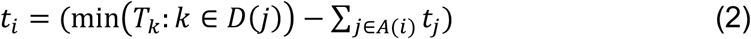

The length of the terminal edges is determined by the length of the ancestral interior edges and the overall length constraint:

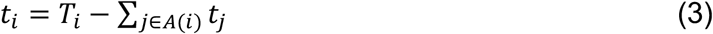

rtreefit assumes that there are *p* − 1 change points in the tree, which correspond to the acquisition of driver mutations. This results in *p* mutation rates *λ*_*j*_ that apply throughout the tree. rtreefit allows for at most one change point each branch and the initial ancestral or wild-type rate is *λ*_0_. Additional rate change points occur at a fraction *α*_*j*_ along branch *j* and descendent branches have the rate *λ*_*j*_ unless there are additional change points in the descendant branches. The effective rate on branches with a change point going from *λ*_*l*_ to *λ*_*j*_ is just the weighted average *α*_*j*_*λ*_*l*_ + (1 − *α*_*j*_)*λ*_*j*_ where we use a uniform, unit-interval prior for the *α* values.

rtreefit assumes that the underlying mutation process follows a Poisson distribution with piecewise-constant, driver specific mutation rates. The number of observed mutations observed on branch *i* in time *t*_*i*_ measured in years: *m*_*i*_ *∼* Poisson(*λt*_*i*_*S*_*i*_) where *S*_*i*_ *∼* Beta 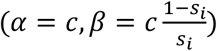 where the parameter *c* = 100. In addition, 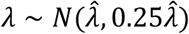 where 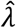 is the naive estimation of a single rate *λ* as the per-patient median of the ratio of the root-to-tip mutation count and the tip-sampling age, and finally rtreefit uses the informative prior for the stick breaking fractions:

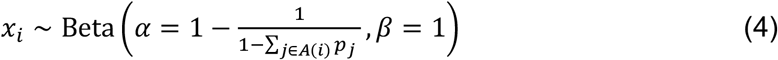

where the *p*_*i*_ is an initial approximation of the duration of the branch length expressed as a fraction of the sampling time:

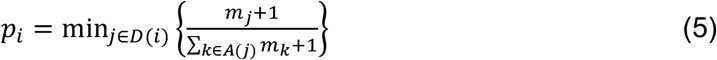

rtreefit considering the higher mutation burden during development by adding a time-dependent excess mutation rate and fitting a logistic function to the mutation burden at three different ages^49,65^:

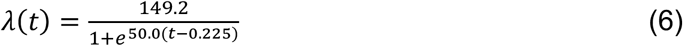

Initially, this maintains an excess instantaneous mutation rate of approximately 150 yr^−1^ before rapidly dropping to zero between two and four months and integrating to an excess of 33.5 mutations by six months – when added to the steady state rate of 18 mutations per year this implies a mutation burden at birth of around 50 mutations.

### Tissue-level phylogenetic reconstruction

Only SSNVs were used to infer the phylogenetic topology using a neighbor-joining method implemented in Biopython, with the reference sequence without any somatic mutations as the root. But both SSNVs and indels were subsequently assigned to the branches of phylogenetic trees. Variants with zero genotype in genomic regions identified as carrying a chromosomal LOH or deletion were set to missing. Since *APC* loss of heterozygosity is a critical event in CRC development, all 5q LOH events and crucial non-silent SSNVs/indels were labelled on the phylogenetic tree.

### Quantify the clonality score of tissues

To assessment of the cell origin over the tissue area, we defined a novel evaluation clonality index - clonality score (CS). Leveraging somatic mutation accumulation within the raw trees, we directly defined the relative length of each clade and calculated the log-transfomed CS as a function of the clonality of the whole tissue as follow:

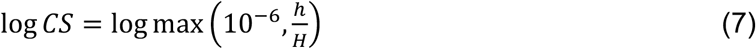

Where *h* represents the relative length of trunk clade of the tree; and *H* is the overall height of the whole tree. The logarithmic transformation ensures a minimal score. Integrating the context of the complete phylogenetic tree and the site frequency spectrum, a CS threshold of 0.05 was applied to distinguish monoclonal tissues from polyclonal tissues.

### Classification of spatial lineage segregation and intermixing

We classified the spatial pattern of each tissue lesion as either intermixing or segregation, based on genetic similarity between crypts across anatomical regions. Single crypts resembling normal tissue on the phylogenetic tree were excluded from the analysis. Lesions were selected according to the following criteria: 1) a minimum of eight crypts distributed across at least three regions, with at least two regions containing two or more crypts each; or 2) a minimum of four regions, with at least three regions containing two or more crypts each. For each tissue, all possible combinations of regions were considered, including intra- and inter-region same-region comparisons (**Supplementary Fig. 8a**). For each crypt pair, similarity *s* was calculated as the proportion of consistent mutation states according to their binary mutation profiles:

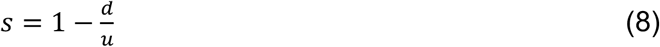

where *d* is the number of different mutation states between the two crypts and *u* is the number of sites mutated in at least one of the two crypts. Regional similarity scores were determined following 10 repeated sampling iterations. These scores range from 0 to 1, with higher values indicating greater genetic similarity. A lesion was classified as intermixing if its maximum inter-region similarity was above the 75th percentile or its inter-to-intra similarity ratio exceeded the median. Otherwise, it was classified as segregating. This classification was then refined using phylogeographic information.

### 2D lattice simulation of tumor growth

We developed a two-dimensional stochastic cellular automaton model to simulate spatial tumor evolution, focusing on the interaction between non-canonical AKP (NC) mutations and an APC mutation. The tissue is represented as a discrete square lattice

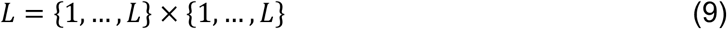

where each lattice site *x* ∈ *L* can be occupied by at most one cell.

Each cell *i* is characterized by a state vector

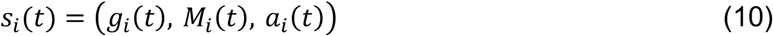

where *g*_*i*_ ∈ {*WT, NC, APC*} denotes the mutation genotype, *M*_*i*_ records the mutation history, and *a*_*i*_ denotes the generation number.

For each lattice site *x*, we define a local neighborhood

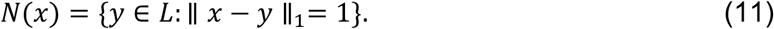

corresponding to nearest-neighbor interactions. Let

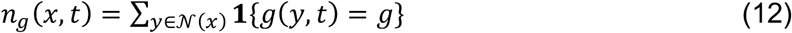

denote the number of neighboring cells of genotype *g*.

We define a neighborhood-dependent promotion factor

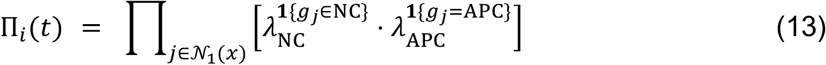

Where *λ*_*NC*_ > 1 is the promotion factor contributed by each neighboring NC-mutant cell; *λ*_*APC*_ > 1 is the promotion factor contributed by each neighboring APC-mutant cell.

At each discrete time step, all cells are updated asynchronously according to the following stochastic processes:

1. Renewal: A cell at site *x* undergoes self-renewal with probability

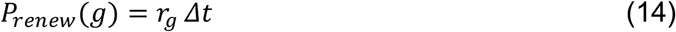

where *r*_*g*_ is the renewal rate associated with genotype *g*. Renewal replaces the cell with a daughter cell at the same location.
2. Proliferation (birth): For a cell *i*, let *E*_1_(*x*) ⊂ *N*_1_(*x*)denote the set of empty neighboring sites. If ∣*E*_1_(*x*)∣ > 0, cell *i* attempts proliferation with probability

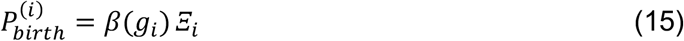

where the genotype-dependent baseline birth rate is

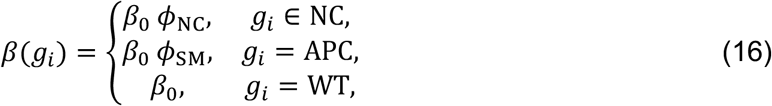

and *Ξ*_*i*_ is a cooperation term:

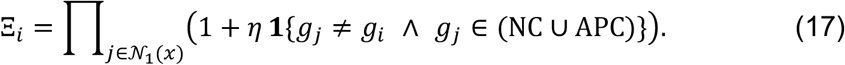

This multiplicative structure ensures that heterotypic mutant neighbors synergistically enhance local proliferation, consistent with non-cell-autonomous fitness effects.
3. Immigration: Cells may also proliferate into second-order neighborhoods *E*_2_(*x*), modeling rare dispersal events. The immigration probability is

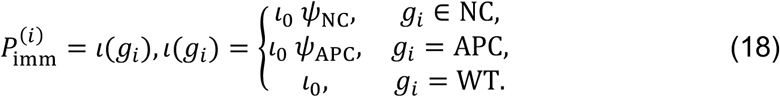

Successful immigration creates a daughter cell at a randomly selected empty site in *E*_2_(*x*).
4. Mutation: During renewal, birth, or immigration, a cell may acquire a new mutation. The mutation probability is modeled as

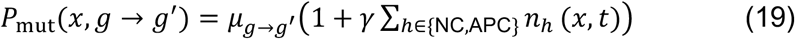

where 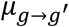 is the baseline mutation rate and *γ* quantifies neighborhood-induced mutagenesis.
5. Death: Cells are removed from the lattice with probability

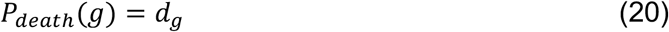

freeing the site for subsequent occupation.
6. Killing (competitive suppression): Cells may attempt to eliminate neighboring mutant cells. For a focal wild-type cell *i*, we define a candidate set

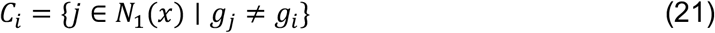

Each candidate *j* is assigned a fitness

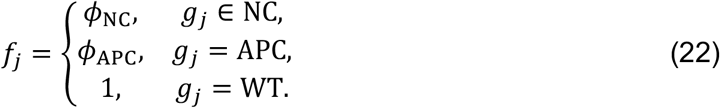

The probability of selecting target *j* is

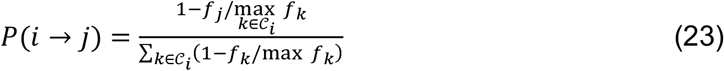

Conditional on selection, the killing attempt succeeds with probability

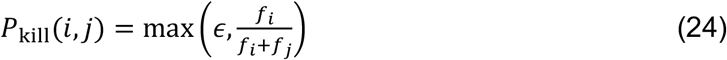

where *ϵ* > 0 enforces a minimal success probability. Successful killing removes the target cell and frees the lattice site.

## Acknowledgments

The authors thank B. Chen for the guidance of single crypt isolations and Hu laboratory members for constructive discussions. This work was supported by National Natural Sciences Foundation of China (32525022, 82241236, 32270693 to Z. Hu), National Key R&D Program of China (2022YFA1304000 to Z.He) and Guangdong S&T Program (2024B1111150001 to Z.He).

## Author contributions

Z. Hu, S.M. and Z. He conceived and designed the study. S.M., H.Z. and L.F. conducted single crypt isolation and sequencing library preparation. S.M. and D.X. analyzed the WGS and WES data. D.X. conducted phylogenetic analysis and inference. K.W. conducted computational simulations. Z.He, J.Y. and J.H. identified patients with synchronous polyps and CRC, performed colonoscopic polypectomy and collected clinical specimens. J.T. conducted pathological analysis. L.T.O.L. provided guidance on experiments. D.D., Z.L., and W.Z. provided guidance on data analysis. S.M., Z.H. and D.X. wrote the manuscript with contributions from all co-authors. Z. Hu and Z. He supervised the project.

## Declaration of Interests

The authors declare no competing interests.

## Data availability

The raw single-crypt WGS data have been deposited in the NGDC’s Genome Sequence Archive (GSA) under accession number HRA010807 (shared link for editor and referees: https://ngdc.cncb.ac.cn/gsa-human/s/d9qCF8dP; the raw sequencing data will be publicly available upon acceptance), HRA009036 (https://ngdc.cncb.ac.cn/gsa-human/browse/HRA009036) and HRA006900 (https://ngdc.cncb.ac.cn/gsa-human/browse/HRA006900).

## Code availability

Source code to generate the figures are available at GitHub (https://github.com/moshl/Single_Crypt_Precancer) and the repository will be made public upon acceptance.

## Supplementary Information

Supplementary Figures 1-11

Supplementary Tables 1-5

---

**Supplementary Figure 1.**
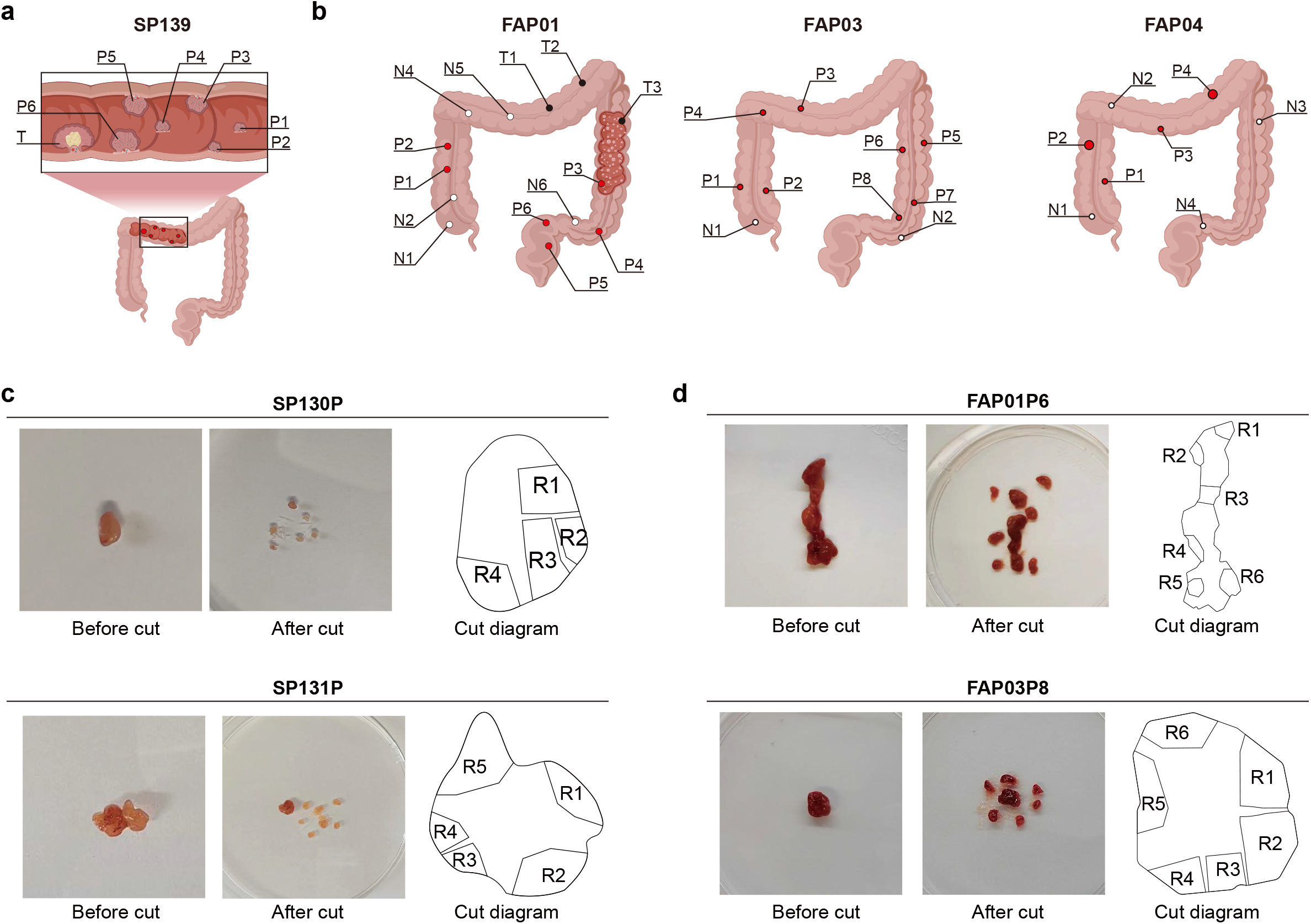
Spatial distribution of multi-regional sampling by anatomical location. **a-b**, Summary schematic showing multi-tissues sampling modalities employed for one sporadic (**a**) and three FAP patients (**b**). **c-d**, Representative images of two sporadic polyps (**c**) and two hereditary polyps (**d**) showing tissue morphology before and after cutting, with corresponding schematic diagrams (right panel) illustrating the multi-region sampling strategy and the boundaries of isolated glandular regions subjected to single-crypt whole-genome sequencing. The colon in **a** and **b** was adapted from the licensed images (License ID: OAYYP4f3bb) provided by Figdraw (https://www.figdraw.com/).

**Supplementary Figure 2.**
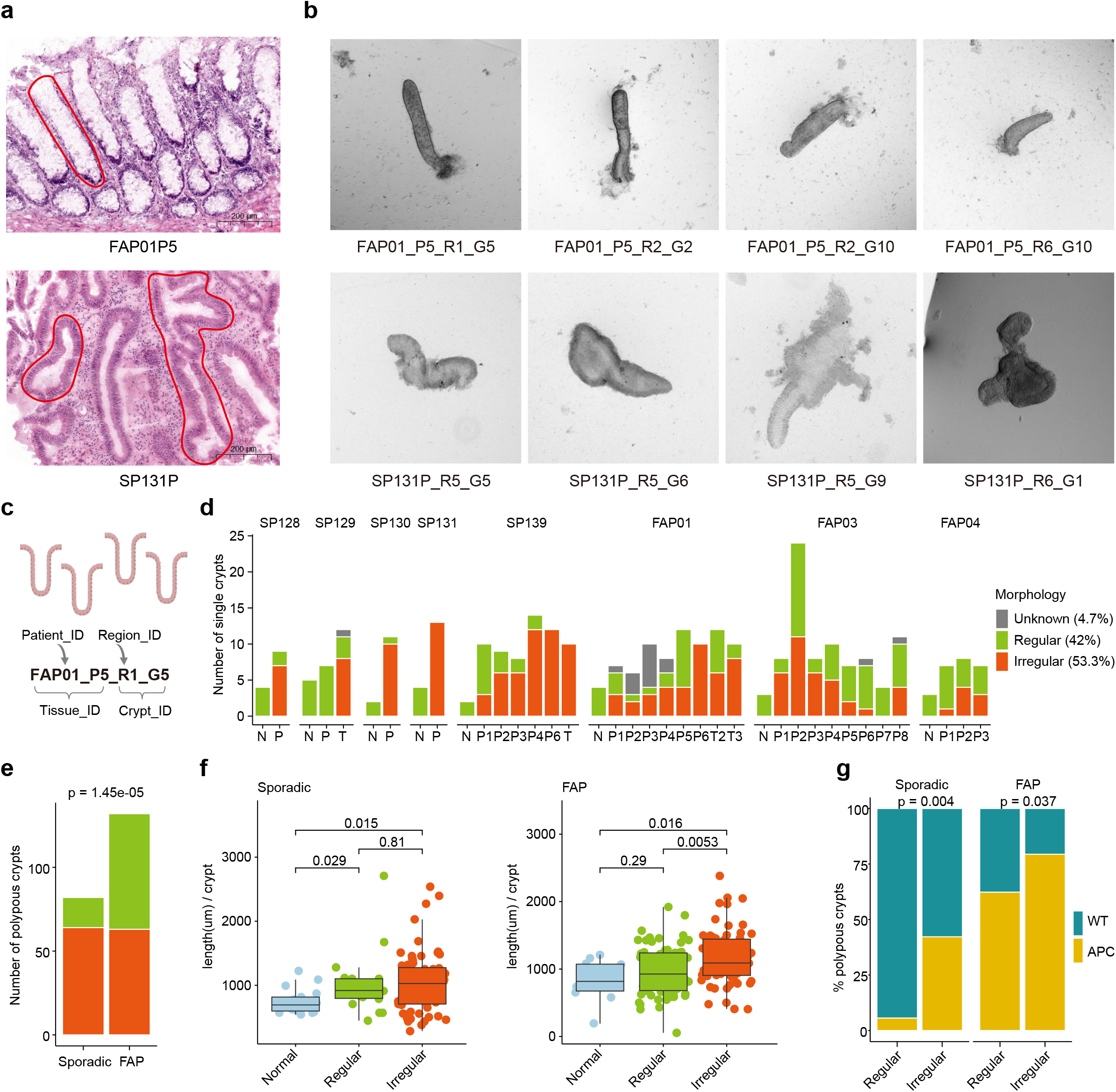
Morphological assessment of crypts within colorectal polyps. **a**, Representative images of haematoxylin and eosin (H&E) staining for tissue section. Red lines outline crypt architectures: representative regular crypts in tissue FAP01P5 and irregular crypts in tissue SP131P. Irregular crypts usually display shape distortion and crypt fusion. Scale bar, 200um. **b**, Representative micrographs of manually isolated crypts corresponding to the regions shown in a, illustrating regular and irregular morphology under microscopy. Crypt barcode assignment is illustrated in **c. c**, Illustrates how a gland is collected and assigned a barcode at each step. Individual crypts were isolated for qualitative morphological examination. Starting from the disease subtype (SP for sporadic or FAP) and the dornor (01), the tissue (P5), region (R1), and individual crypts (or glands, labeled as G) are assigned respectively. **d**, Distribution of crypt morphological types (regular, irregular and unknown labeled by different colors) across all tissues for each patient. **e**, Comparison of regular versus irregular crypt counts between sporadic and hereditary (FAP) polyps. *P* value by two-sided Fisher’s exact test. **f**, Comparison of length across normal crypts, polyp-derived regular/irregular crypts. In box plots, the horizontal line is the median, the box delineates the 25th to 75th centiles, and whiskers extend to 1.5 times the interquartile range. *P* value by two-sided Wilcoxon rank-sum tests. **g**, Comparison of the proportion of APC-mutant crypts across regular and irregular crypt architecture within polyps from sporadic and FAP (right).

**Supplementary Figure 3.**
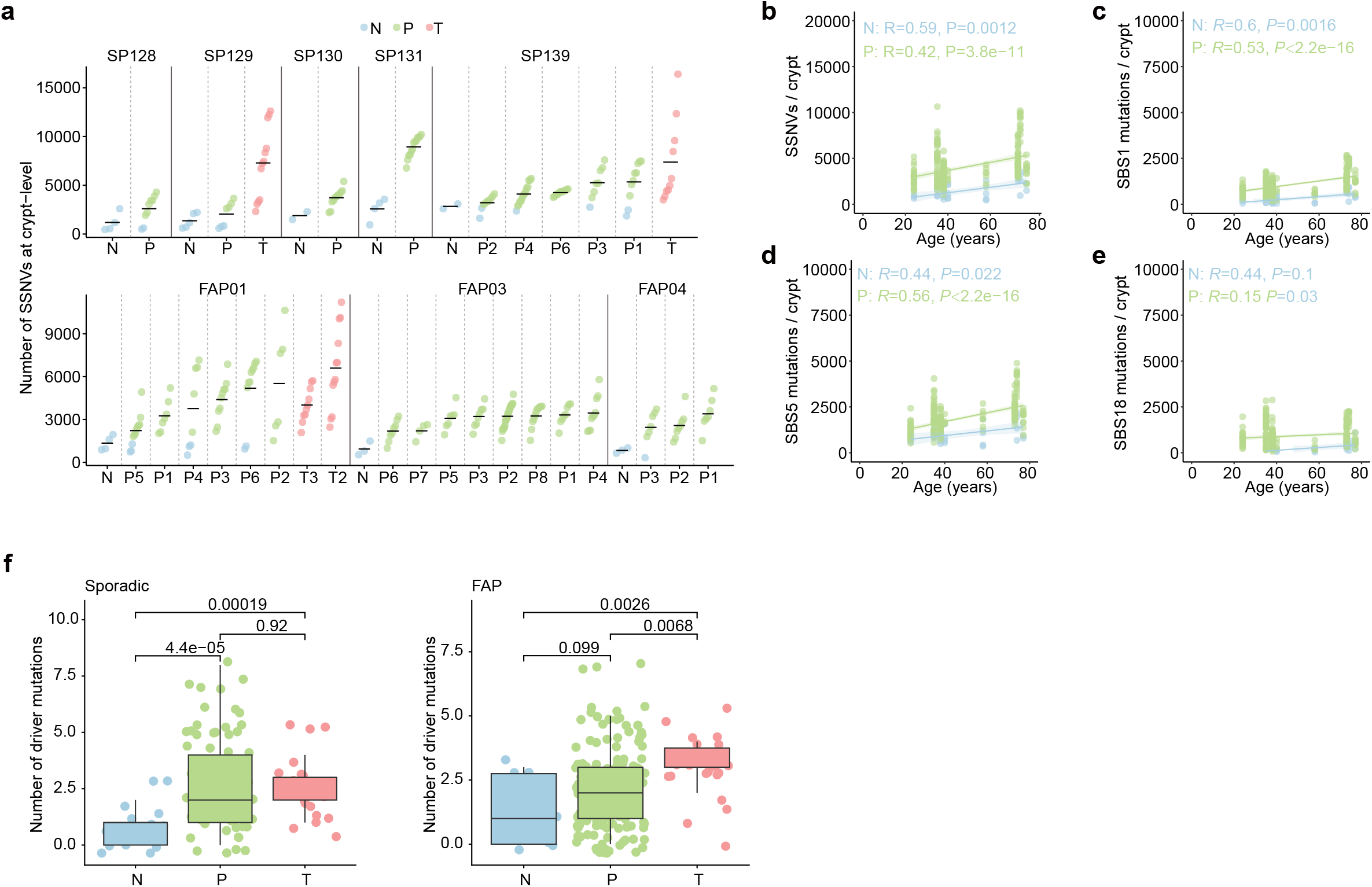
Somatic single nucleotide variants (SSNVs) in normal, polyps and tumor crypts. **a**, Number of SSNVs detected in each crypt. Each point represents a single crypt, colored by tissues. Black horizontal lines denote medians. **b-e**, Correlation analyses between donor age and mutation burdens at the crypt level. Total mutation burden (**b**) and the burdens of ubiquitous mutational signatures SBS1 (**c**), SBS5 (**d**), and SBS18 (**e**) were plotted against age for normal mucosa and polyps separately. Pearson’s *r* and *P* value are shown. **f**, Comparison of somatic driver mutations (**Methods**) across normal, polyp and tumor crypts. *P*-value by two-sided Wilcoxon rank-sum tests.

**Supplementary Figure 4.**
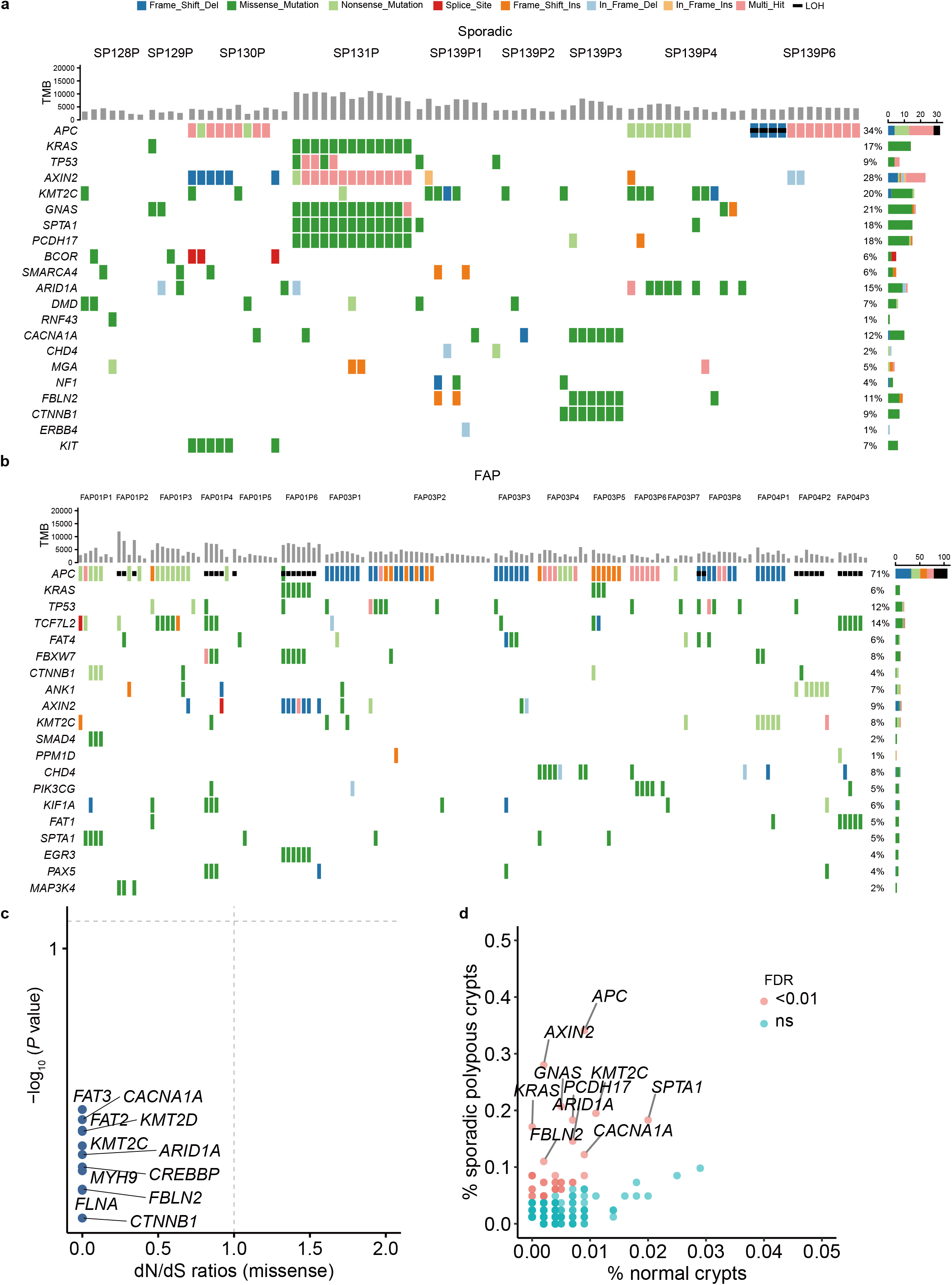
CRC driver mutational landscape of crypts during polyp initiation. **a-b**, Onco-plots summarizing the landscape of non-silent SSNVs, Indels and copy-neutral LOH (LOH) in CRC driver genes based on single-crypt WGS spanning normal mucosae, premalignant polyps and synchronous tumors from sporadic (**a**) and FAP (**b**) patients. Only somatic alterations are shown. **c**. dN/dS ratios of enriched genes in **Fig. 2e** for missense mutations in normal crypts from sporadic patients. *P* values for the log likelihood-ratio test are shown. Vertical and horizontal lines correspond to absolute dN/dS values of 1 and *P* < 0.1. **d**. Prevalence of the CRC-associated drivers in normal crypts (Lee-Six et al 2019, n = 557) versus sporadic polyp crypts (n = 94). *P* value was calculated using two-sided Fisher’s exact test and adjusted by Benjamini-Hochberg method (FDR < 0.01). Marked genes have FDR < 0.01 and prevalence ≥ 10% in sporadic derived crypts.

**Supplementary Figure 5.**
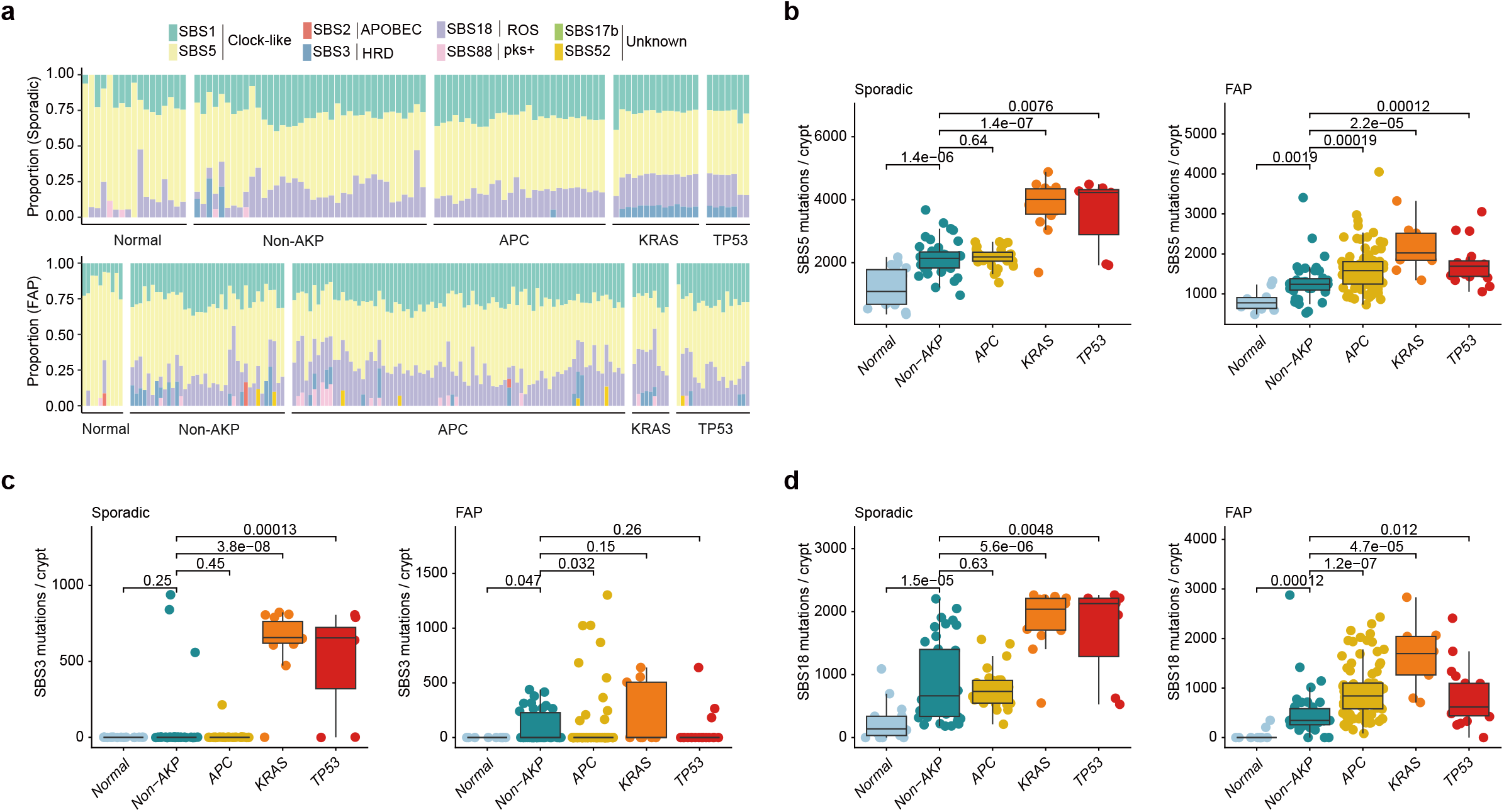
Mutational Signatures across normal, non-AKP and AKP crypts. **a**. Substitution mutational signatures for normal and adenoma single crypts in sporadic and FAP patients. **b-d**. Comparison of SBS5 (**b**), SBS3 (**c**) and SBS18 (**d**) burdens across normal, non-AKP and AKP crypts from sporadic (left) and FAP (right) patients, respectively. *P*-value by two-sided Wilcoxon rank-sum test.

**Supplementary Figure 6.**
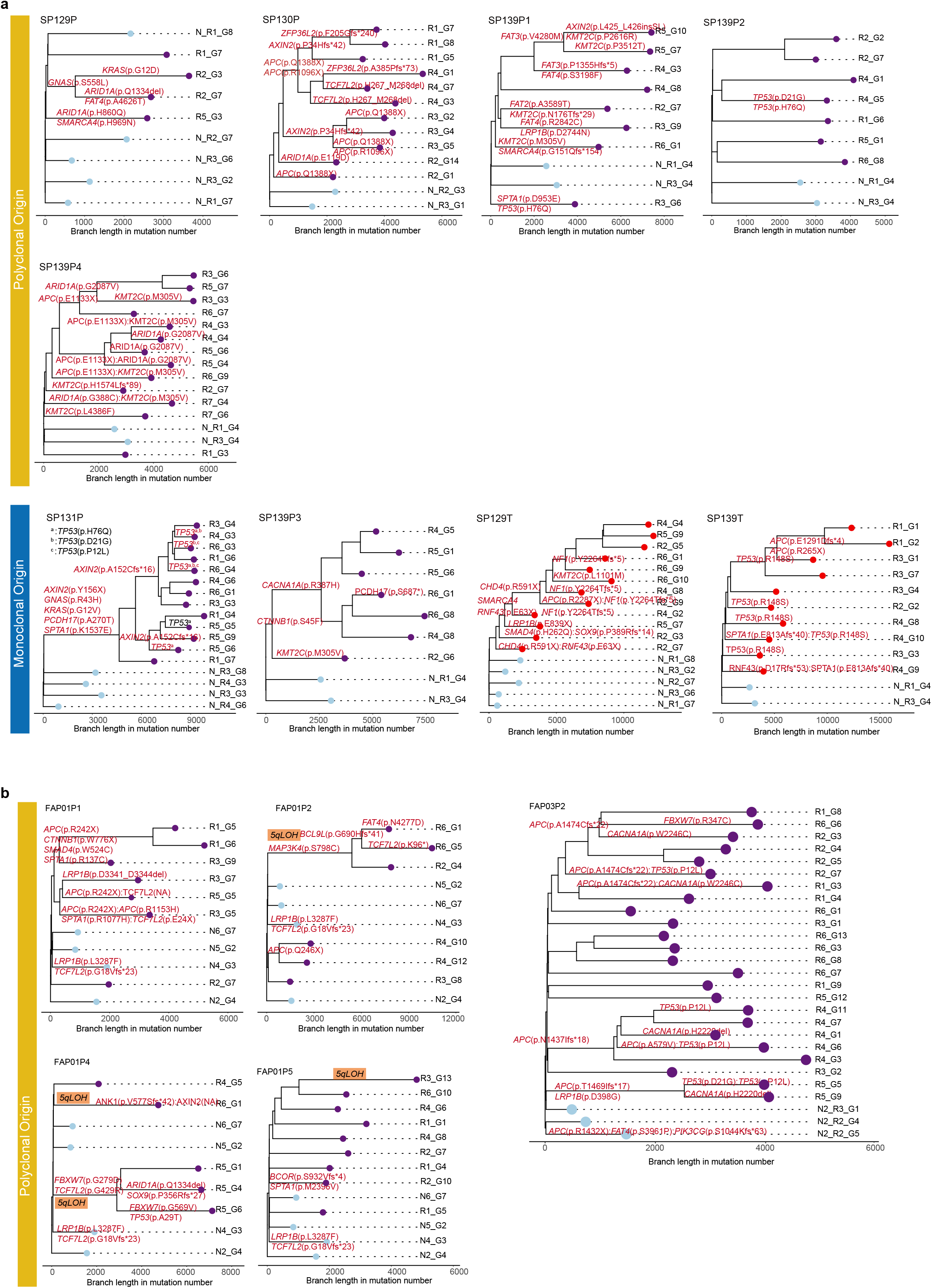
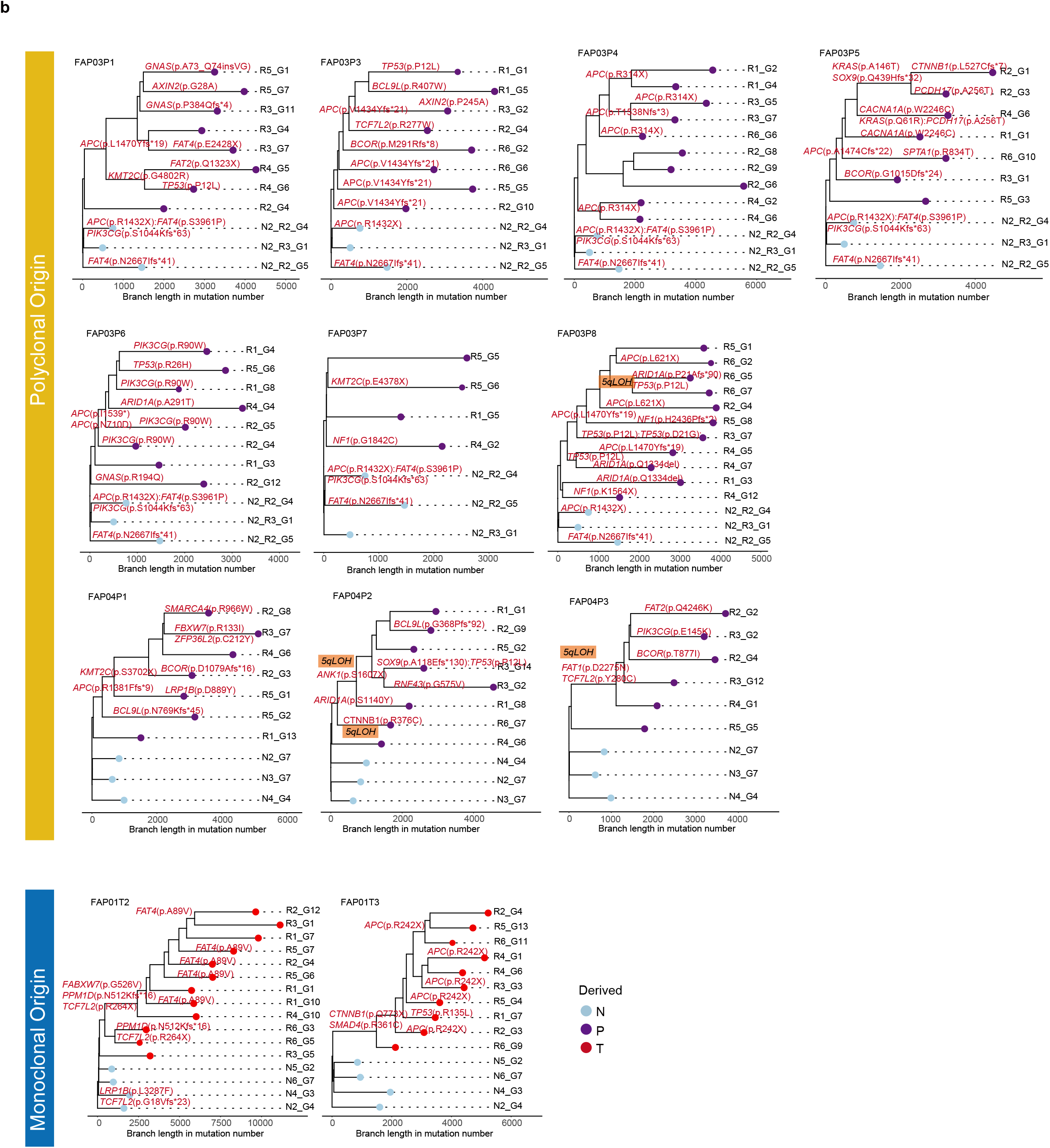
Phylogenetic tree showing the clonality of sporadic and hereditary tissues. **a-b**, Single-crypt phylogenies including neoplastic crypts from an individual polyp and normal intestinal crypts from sporadic (**a**) and FAP (**b**) patients. Tips represent individual crypts. Preneoplastic/Neoplastic crypts are colored in purple/red while normal colon crypts from corresponding adjacent colons are colored in light blue. Key driver somatic mutations and CNV events are annotated.

**Supplementary Figure 7.**
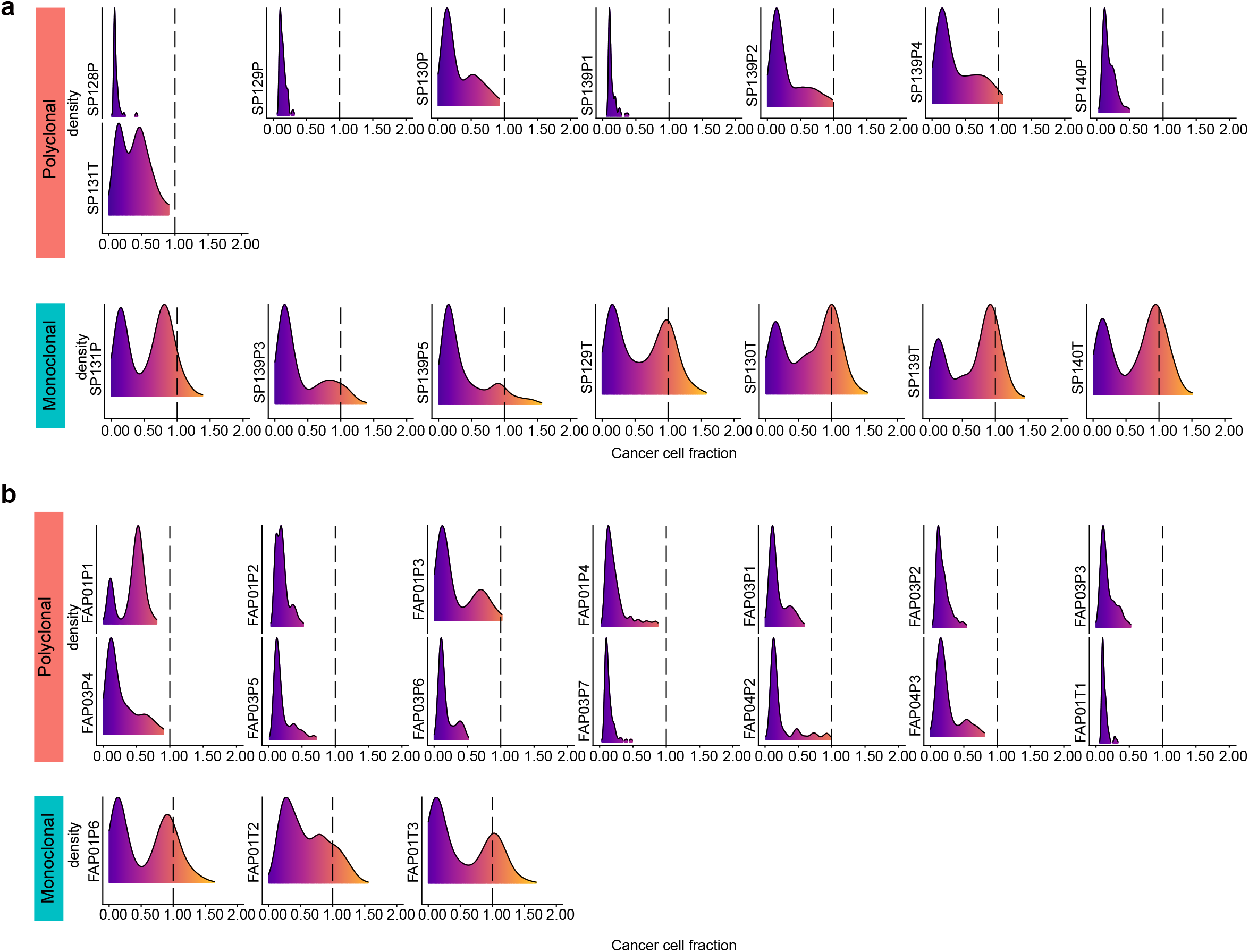
The site frequency spectrum of polyclonal versus monoclonal tissues. **a**, The CCF distributions of SSNVs/Indels for 8 polyclonal tissues and 7 monoclonal tissues from sporadic patients. **b**, The CCF distributions of SSNVs/Indels for 14 polyclonal tissues and 3 monoclonal tissues from FAP patients. Tissues of monoclonal origin are characterized by a clonal mutational cluster centered around CCF=1.0, whereas CCFs of polyclonal tissues are all below 1.

**Supplementary Figure 8.**
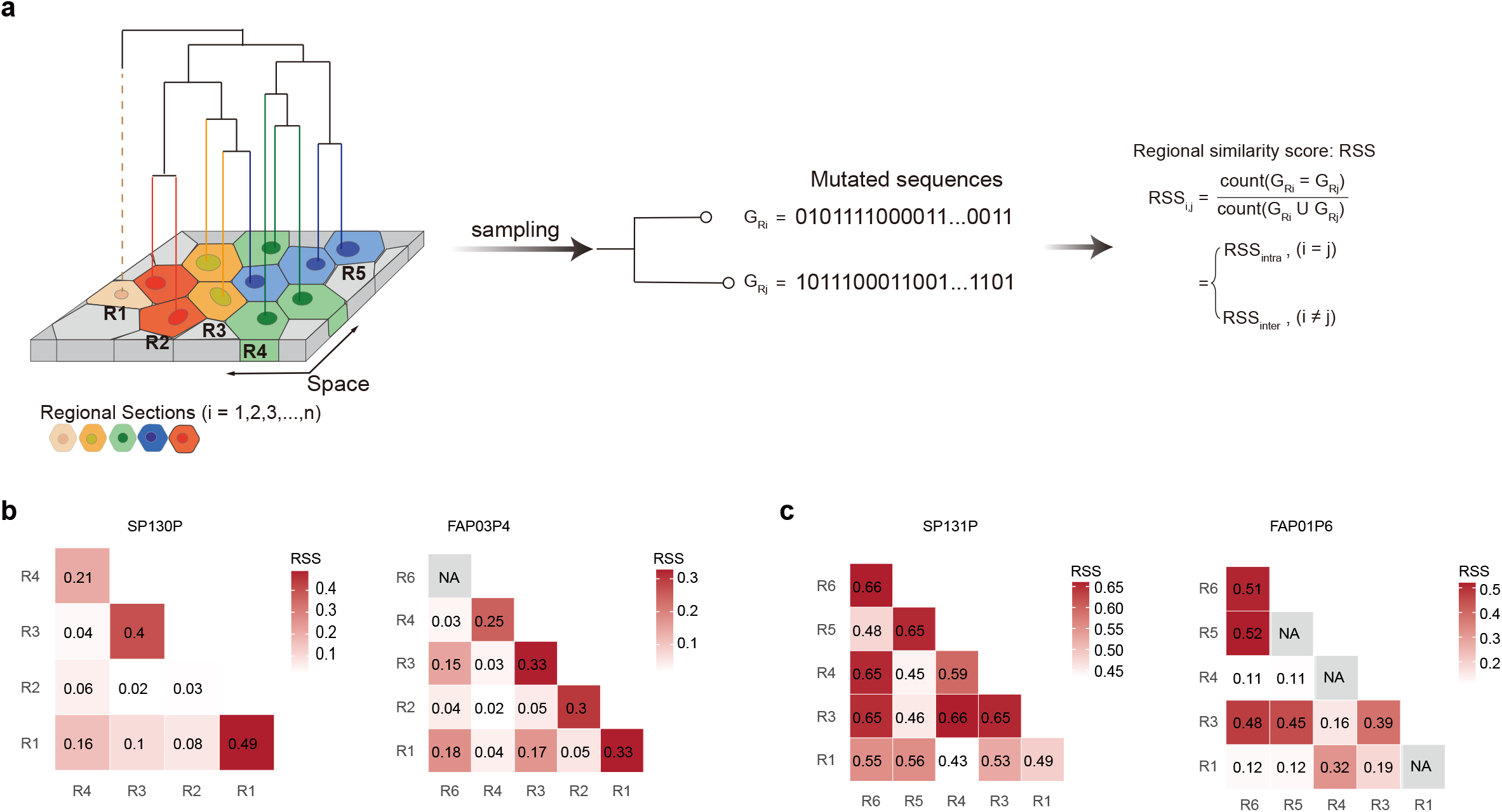
Tissue-level phylogenetic trees of multi-regional crypts and inter/intra-polyp similarities. **a**, Schematic of the spatial heterogeneity of polyps or tumors. **b-c**, Heatmaps showing the pairwise mutational similarity of random crypt pairs within a region or between regions from the same polyps with spatial-segregation (**b**) and spatial-mixing (**c**).

**Supplementary Figure 9.**
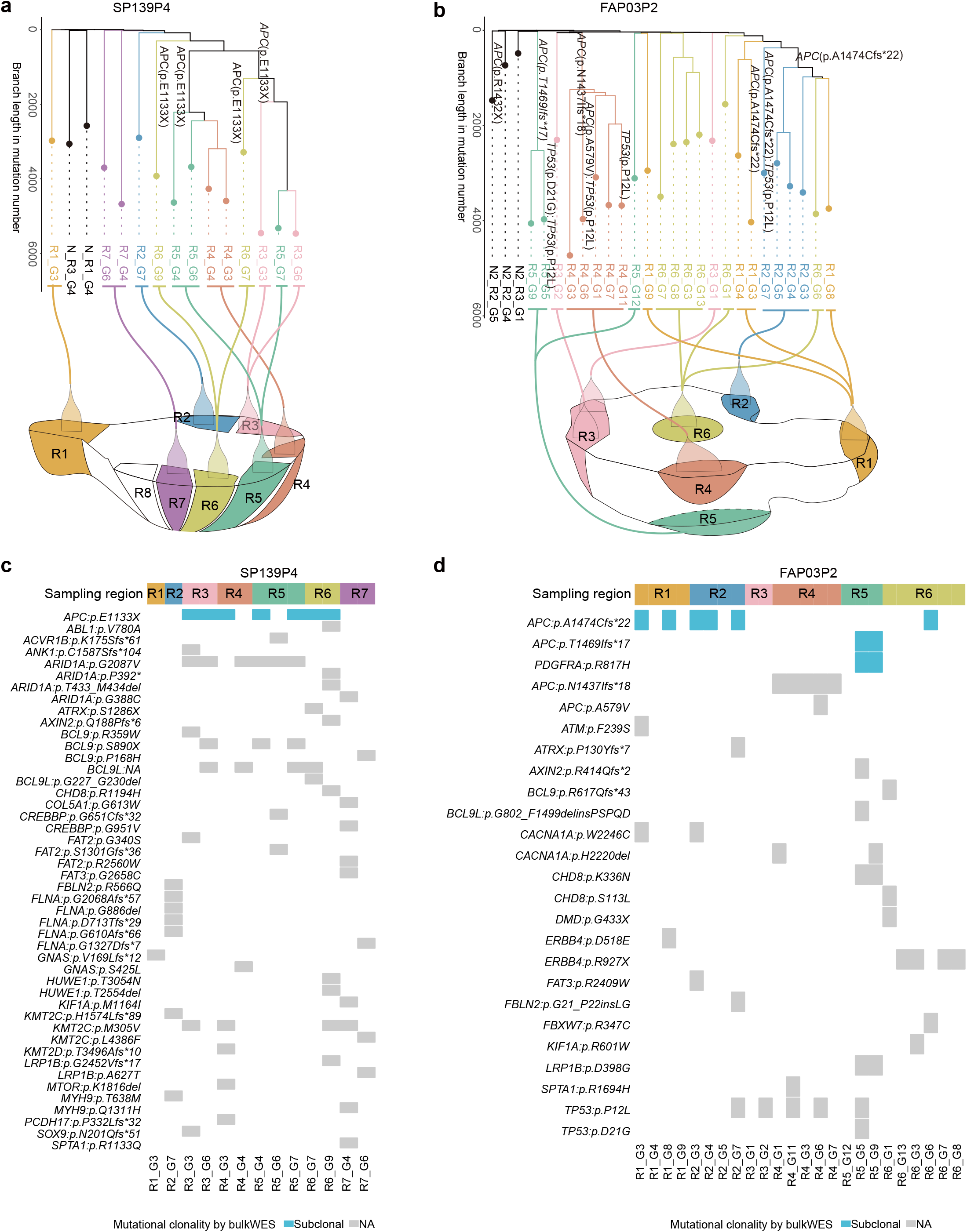
Phylogeography reveals the spatial imbalanced clonal expansion. **a-b**, Single-crypt phylogeny (top) and corresponding sampling regions (bottom) for a representative sporadic (**a**) and FAP (**b**) polyp. **c-d**, Heatmaps indicated the presence of representative subclonal mutations in putative CRC drivers across multiple crypts per polyp, where tissue-level bulk whole-exome sequencing of the matched polyp is included for comparison.

**Supplementary Figure 10.**
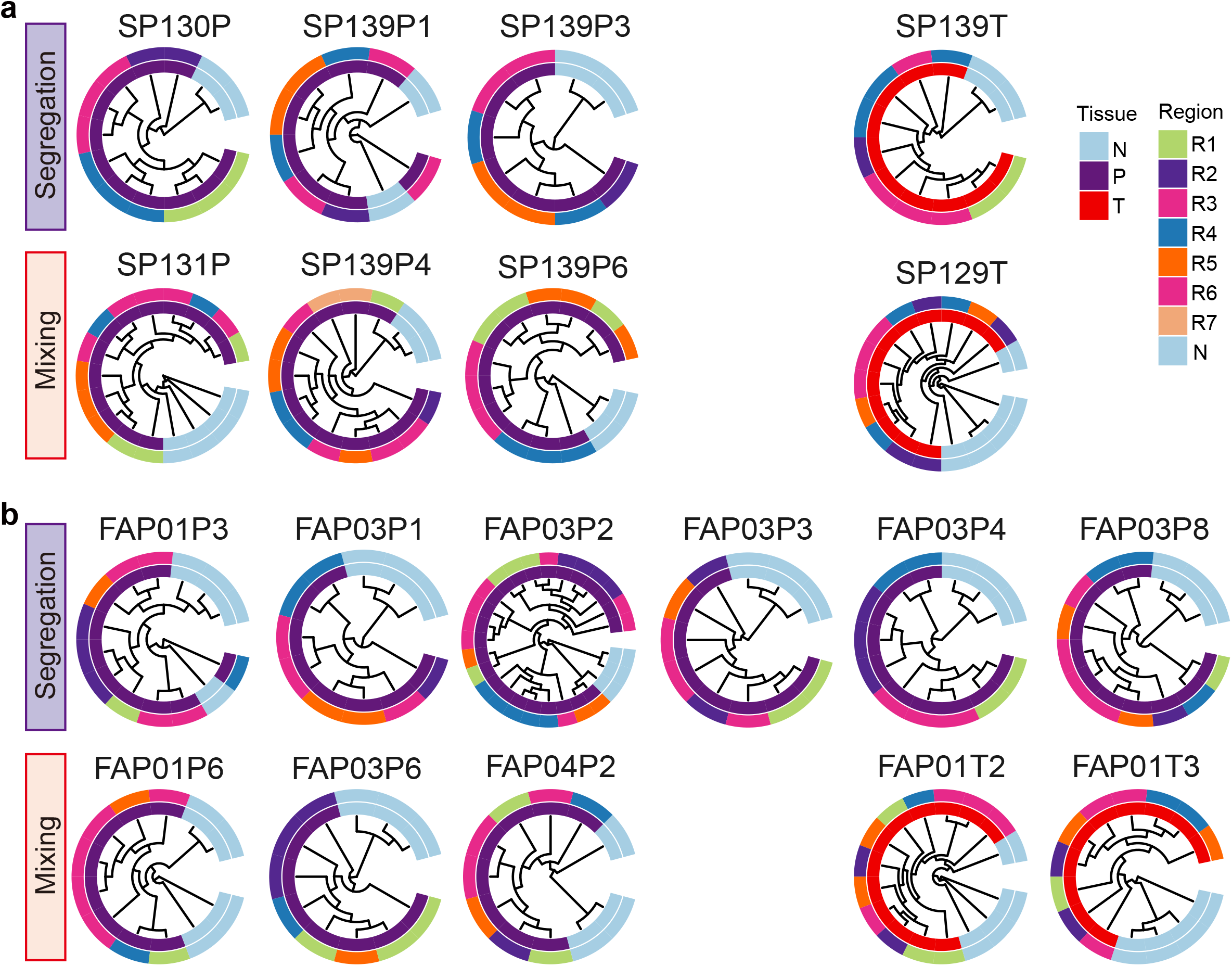
Tissue-level phylogenetic trees of single-crypt from multiple regions within polyps or tumors. **a-b**, Phylogenetic trees of single crypts from multiple polyps, tumors and adjacent normal tissues with spatial mixing mode or spatial segregation mode from all sporadic (**a**) and FAP (**b**) patients.

**Supplementary Figure 11.**
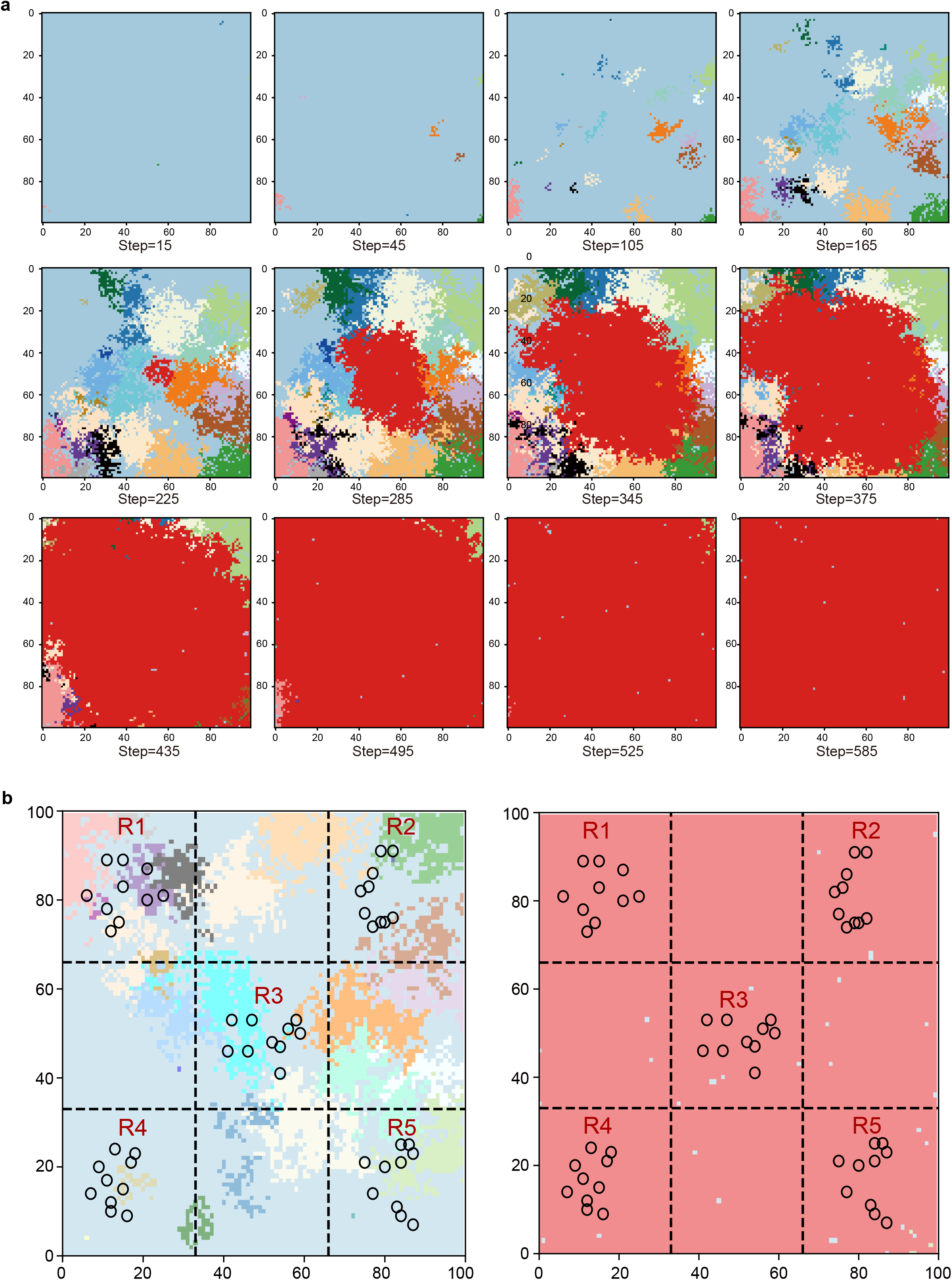
Spatial tumor growth simulation and region-based lineage sampling. **a**, Representative snapshots of the two-dimensional lattice-based tumor growth simulation at 12 selected time points, illustrating the spatial expansion and clonal diversification of tumor cells over time. Each panel shows the full 100×100 lattice, with colors indicating distinct clones. **b**, Schematic of the spatial sampling strategy used to construct lineage trees from the simulated tumor. At each time point, the lattice was partitioned into a 3×3 grid of equally sized regions, from which five spatially separated regions were selected: the four corner regions and the central region, denoted as R1-R5. Within each region, 10 cells were randomly sampled, yielding a total of 50 cells per time point for lineage reconstruction. The spatial locations of sampled regions were fixed across time points, allowing comparison of lineage tree structures during polyclonal and monoclonal growth phases.

## Notes

### Competing Interest Statement

The authors have declared no competing interest.

## References

1 Spira, A. et al. Precancer Atlas to Drive Precision Prevention Trials. Cancer Res 77, 1510–1541 (2017).

2 Derks, L. L. M. & van Boxtel, R. Stem cell mutations, associated cancer risk, and consequences for regenerative medicine. Cell Stem Cell 30, 1421–1433 (2023).

3 Faupel-Badger, J. et al. Defining precancer: a grand challenge for the cancer community.Nat Rev Cancer 24, 792–809 (2024).

4 Stangis, M. M. et al. The Hallmarks of Precancer. Cancer Discov 14, 683–689 (2024).

5 Zhang, S. et al. Tumor initiation and early tumorigenesis: molecular mechanisms and interventional targets. Signal Transduct Target Ther 9, 149 (2024).

6 Weeden, C. E., Hill, W., Lim, E. L., Gronroos, E. & Swanton, C. Impact of risk factors on early cancer evolution. Cell 186, 1541–1563 (2023).

7 Gerstung, M. et al. The evolutionary history of 2,658 cancers. Nature 578, 122–128 (2020).

8 Leshchiner, I. et al. Inferring early genetic progression in cancers with unobtainable premalignant disease. Nat Cancer 4, 550–563 (2023).

9 Curtius, K., Wright, N. A. & Graham, T. A. Evolution of Premalignant Disease. Cold Spring Harb Perspect Med 7 (2017).

10 Yang, J. et al. Signaling pathways and targeted interventions for precancers. Signal Transduct Target Ther 11, 9 (2026).

11 Kuipers, E. J. et al. Colorectal cancer. Nat Rev Dis Primers 1, 15065 (2015).

12 Cancer Genome Atlas, N. Comprehensive molecular characterization of human colon and rectal cancer. Nature 487, 330–337 (2012).

13 Fearon, E. R. & Vogelstein, B. A genetic model for colorectal tumorigenesis. Cell 61, 759–767 (1990).

14 Groden, J. et al. Identification and characterization of the familial adenomatous polyposis coli gene. Cell 66, 589–600 (1991).

15 Galiatsatos, P. & Foulkes, W. D. Familial adenomatous polyposis. Am J Gastroenterol 101, 385–398 (2006).

16 Becker, W. R. et al. Single-cell analyses define a continuum of cell state and composition changes in the malignant transformation of polyps to colorectal cancer. Nat Genet 54, 985–995 (2022).

17 Chen, B. et al. Differential pre-malignant programs and microenvironment chart distinct paths to malignancy in human colorectal polyps. Cell 184, 6262–6280 e6226 (2021).

18 Van Egeren, D. et al. Polyclonal origins of human premalignant colorectal lesions. Nature 650, 1017–1024 (2026).

19 Lu, Z. et al. Polyclonal-to-monoclonal transition in colorectal precancerous evolution.Nature 636, 233–240 (2024).

20 Sadien, I. D. et al. Polyclonality overcomes fitness barriers in Apc-driven tumorigenesis.Nature 634, 1196–1203 (2024).

21 Gaynor, L. et al. Crypt density and recruited enhancers underlie intestinal tumour initiation. Nature 640, 231–239 (2025).

22 Islam, M. et al. Temporal recording of mammalian development and precancer. Nature 634, 1187–1195 (2024).

23 Nowell, P. C. The clonal evolution of tumor cell populations. Science 194, 23–28 (1976).

24 Kuiken, M. C., Witsen, M., Voest, E. E. & Dijkstra, K. K. Strength through diversity: how cancers thrive when clones cooperate. Mol Oncol 20, 226–247 (2026).

25 Potten, C. S., Kellett, M., Roberts, S. A., Rew, D. A. & Wilson, G. D. Measurement of in vivo proliferation in human colorectal mucosa using bromodeoxyuridine. Gut 33, 71–78 (1992).

26 Gehart, H. & Clevers, H. Tales from the crypt: new insights into intestinal stem cells. Nat Rev Gastroenterol Hepatol 16, 19–34 (2019).

27 Lee-Six, H. et al. The landscape of somatic mutation in normal colorectal epithelial cells. Nature 574, 532–537 (2019).

28 Cagan, A. et al. Somatic mutation rates scale with lifespan across mammals. Nature 604, 517–524 (2022).

29 Olafsson, S. et al. Somatic Evolution in Non-neoplastic IBD-Affected Colon. Cell 182, 672–684 e611 (2020).

30 Heide, T. et al. The co-evolution of the genome and epigenome in colorectal cancer. Nature 611, 733–743 (2022).

31 Sottoriva, A. et al. A Big Bang model of human colorectal tumor growth. Nat Genet 47, 209–216 (2015).

32 Househam, J. et al. Phenotypic plasticity and genetic control in colorectal cancer evolution. Nature 611, 744–753 (2022).

33 Ellis, P. et al. Reliable detection of somatic mutations in solid tissues by laser-capture microdissection and low-input DNA sequencing. Nat Protoc 16, 841–871 (2021).

34 Xie, D. et al. Deciphering genomic evolution of metastatic organotropism with 535 paired primary lung cancers and metastases. Cell Rep 44, 116449 (2025).

35 Hu, Z., Li, Z., Ma, Z. & Curtis, C. Multi-cancer analysis of clonality and the timing of systemic spread in paired primary tumors and metastases. Nat Genet 52, 701–708 (2020).

36 Koh, G., Degasperi, A., Zou, X., Momen, S. & Nik-Zainal, S. Mutational signatures: emerging concepts, caveats and clinical applications. Nat Rev Cancer 21, 619–637 (2021).

37 Li, Y. et al. Patterns of somatic structural variation in human cancer genomes. Nature 578, 112–121 (2020).

38 Gori, K. & Baez-Ortega, A. sigfit: flexible Bayesian inference of mutational signatures. bioRxiv, 372896 (2020).

39 Nik-Zainal, S. et al. Mutational processes molding the genomes of 21 breast cancers. Cell 149, 979–993 (2012).

40 Alexandrov, L. B. et al. Signatures of mutational processes in human cancer. Nature 500, 415–421 (2013).

41 Pleguezuelos-Manzano, C. et al. Mutational signature in colorectal cancer caused by genotoxic pks(+) E. coli. Nature 580, 269–273 (2020).

42 Rodriguez-Martin, B. et al. Pan-cancer analysis of whole genomes identifies driver rearrangements promoted by LINE-1 retrotransposition. Nat Genet 52, 306–319 (2020).

43 Zumalave, S. et al. Concurrent L1 retrotransposition events promote reciprocal translocations in human tumorigenesis. Science, eaee4513 (2026).

44 Nam, C. H. et al. Widespread somatic L1 retrotransposition in normal colorectal epithelium. Nature 617, 540–547 (2023).

45 Chu, C. et al. Comprehensive identification of transposable element insertions using multiple sequencing technologies. Nat Commun 12, 3836 (2021).

46 Rausch, T. et al. DELLY: structural variant discovery by integrated paired-end and split-read analysis. Bioinformatics 28, i333–i339 (2012).

47 Lee, B. C. H. et al. Mutational landscape of normal epithelial cells in Lynch Syndrome patients. Nat Commun 13, 2710 (2022).

48 Williams, N. et al. Life histories of myeloproliferative neoplasms inferred from phylogenies. Nature 602, 162–168 (2022).

49 Spencer Chapman, M. et al. Lineage tracing of human development through somatic mutations. Nature 595, 85–90 (2021).

50 Marvalim, C. & Chan, D. K. H. Early mutational events and clonal dynamics in normal crypts: implications for colorectal tumorigenesis. Hum Genomics 19, 146 (2025).

51 Lourenco, F. C. et al. Decay of driver mutations shapes the landscape of intestinal transformation. Nature 649, 729–738 (2026).

52 Seferbekova, Z., Lomakin, A., Yates, L. R. & Gerstung, M. Spatial biology of cancer evolution. Nat Rev Genet 24, 295–313 (2023).

53 Wang, R. G. et al. Usefulness of analyzing endoscopic features in identifying the colorectal serrated sessile lesions with and without dysplasia. World J Clin Cases 11, 6995–7003 (2023).

54 Orlando, F. A. et al. Aberrant crypt foci as precursors in colorectal cancer progression. J Surg Oncol 98, 207–213 (2008).

55 Fenoglio-Preiser, C. M. & Noffsinger, A. Aberrant crypt foci: A review. Toxicol Pathol 27, 632–642 (1999).

56 Orlowska, J. Serrated polyps of the colorectum: histological classification and clinical significance. Pol J Pathol 61, 8–22 (2010).

57 Zhao, R. & Michor, F. Patterns of proliferative activity in the colonic crypt determine crypt stability and rates of somatic evolution. PLoS Comput Biol 9, e1003082 (2013).

58 McKenna, A. et al. The Genome Analysis Toolkit: A MapReduce framework for analyzing next-generation DNA sequencing data. Genome Res 20, 1297–1303 (2010).

59 Zhao, Q. et al. Comprehensive profiling of 1015 patients exomes reveals genomic-clinical associations in colorectal cancer. Nat Commun 13, 2342 (2022).

60 Ha, G. et al. TITAN: inference of copy number architectures in clonal cell populations from tumor whole-genome sequence data. Genome Res 24, 1881–1893 (2014).

61 Wang, Y. et al. APOBEC mutagenesis is a common process in normal human small intestine. Nat Genet 55, 246–254 (2023).

62 Alexandrov, L. B. et al. The repertoire of mutational signatures in human cancer. Nature 578, 94–101 (2020).

63 Martincorena, I. et al. Universal Patterns of Selection in Cancer and Somatic Tissues.Cell 171, 1029–1041 e1021 (2017).

64 Coorens, T. H. H. et al. Reconstructing phylogenetic trees from genome-wide somatic mutations in clonal samples. Nat Protoc 19, 1866–1886 (2024).

65 Osorio, F. G. et al. Somatic Mutations Reveal Lineage Relationships and Age-Related Mutagenesis in Human Hematopoiesis. Cell Rep 25, 2308–2316 e2304 (2018).

